# Selective removal of green pigments and associated proteins from clover-grass protein concentrates: Molecular insight into a non-destructive, two-stage membrane-based biorefinery concept for high-quality food protein production

**DOI:** 10.1101/2025.10.24.684365

**Authors:** Simon Gregersen Echers, Naim Abdul-Khalek, Nete Hassing Jensen, Anders Kjær Jørgensen, Tuve Mattsson, Mads Koustrup Jørgensen, Peter Stephensen Lübeck, Mette Lübeck

## Abstract

Green leaves are gaining traction as an emerging protein source and a sustainable alternative to animal-based protein as leaf proteins often possess good nutritional and functional properties. Current methods for producing protein isolates from leaves include partially or fully denaturing conditions, diminishing protein solubility and functionality. Here, we characterize the performance of a multi-stage, membrane-based green biorefinery concept capable of producing a native protein product from clover grasses devoid of attributes such as green color and grassy smell/taste. By sampling at each step along the process, we obtain insights on the fate of proteins and pigments over each unit operation. The process efficiently removes green pigments (>99.9%), when comparing the product stream with the initial feed stream, based on UV/Vis analysis. Using mass spectrometry-based proteomics, two complementary quantification strategies, and subsequent bioinformatic analysis, this can particularly be ascribed to a very selective retention (>99%) of unwanted membrane-associated and pigment-binding proteins in the first-stage filtration. In the second-stage filtration and subsequent diafiltration stage, residual unwanted proteins, fragments, and pigments are efficiently washed out while retaining the overall protein composition. The product maintains the high RuBisCO content of green juice and is furthermore enriched in proteins with known antioxidant properties while depleted in known food allergens. This work presents an in-depth understanding of protein-level selectivity in membrane-based green biorefinery and can help guide process optimization towards improved yields and quality.

**Highlights:**

- Streams from a two-stage membrane-based green biorefinery were characterized
- >99.9% pigment-binding proteins were removed primarily in first stage filtration
- The 2^nd^ filtration and diafiltration removed residual pigment and retained protein
- The full process efficiently removed >99.9% of chlorophylls and carotenoids
- The soluble product is rich in RuBisCO and enriched in known antioxidant proteins

## 1. Introduction

In light of the significant environmental impact imposed by conventional livestock and arable land farming, an increasing focus on the need for sustainable food and protein production is becoming evident on a global scale [1,2]. As part of the green transition of society, plants are becoming increasingly important as sources of dietary protein [3,4]. The grand challenge for plant protein-based ingredients in foods is subpar nutritional value and functional properties compared to conventional animal-based protein ingredients. While soy protein to some extent addresses these needs and currently has the largest market share among plant proteins, soy is affiliated with sustainability concerns, emphasizing the need for locally sourced, sustainable alternatives [5]. Among these, leaf protein is generating traction as a future source of food protein not just scientifically but also politically – particularly in Northern Europe [6–9]. Perennial grasses and legumes such as clovers and alfalfa, are gathering increased attention as emerging sources. Not only are they easily cultivated on a global scale, but they also possess multiple benefits such as reduced need for fertilizers and pesticides, drought resistance, improved soil quality, high biomass yields, repeated harvest, and a high protein content (up to 30% of dry matter) [10–16]. Utilization of green leaves may also accommodate a reduction in food waste, as leafy parts of food crops, often constituting upwards of 75% of the biomass from e.g. broccoli and kale, are typically discarded [17].

In photosynthetically active green leaves of plants, such as perennial forage crops like grasses and clovers, the most abundant protein is the enzyme ribulose-1,5-bisphosphate carboxylase/oxygenase (RuBisCO), constituting upwards of 50% of the soluble leaf protein [18,19]. RuBisCO is a hexadecameric globular protein of approximately 550 kDa consisting of eight large (rbcL) and eight small (rbcS) subunits [20]. Because of the high content, not only in leaves but also in marine plants, photosynthetic bacteria, and eukaryotic algae, RuBisCO is considered the most abundant protein on Earth [20]. In addition to the prevalence of the protein, RuBisCO has several advantageous properties in a food context. The amino acid composition makes it highly suitable for human nutrition and furthermore, RuBisCO has been shown to possess very promising attributes as a protein-based food ingredient with e.g. excellent solubility as well as strong foaming and gelling properties [21–25]. Obtaining protein isolates and concentrates with a high content of RuBisCO is therefore considered desirable for producing sustainable protein ingredients for foods.

A major challenge in using leafy greens for protein production is the presence of unwanted constituents, posing not only a problem for functionality but also for sensory attributes and consumer acceptance. One group of unwanted constituents in a protein product is pigments such as chlorophylls and carotenoids. Chlorophylls are highly light sensitive and function as photosensitizers, which in turn may lead to undesired sensory attributes when incorporated in foods [26,27]. While carotenoids are frequently considered beneficial bioactives and used as health-promoting food additives [28], they typically co-localize with other undesired constituents such as chlorophylls [29]. Moreover, other small molecule phytochemicals such as quercetin, a flavonoid polyphenol, and phenolic acids, are associated with lower sensory quality such as bitterness and astringency [30–32]. Avoidance of such undesirable compounds in a protein product requires development of strategies for refining plant leaves in a selective manner.

Most scalable green biorefinery concepts developed thus far are focused on producing feed protein [33–36]. This constitutes a major opportunity for biomass valorization as functional food protein is generally of much higher value than feed protein [37]. The challenge is developing a cost-effective and scalable process [38]. On a more developmental and experimental scale, conventional green biorefinery concepts for producing a “white” food-grade protein from green biomass include initial wet fractionation (typically by screw pressing) to obtain a green juice, followed by a mild heating step (typically around 50°C) to produce a clarified juice and a “green protein” precipitate. The clarified juice is then further processed by precipitating the “white” protein using heating at higher temperatures (typically around 80°C) or isoelectric precipitation under acidic conditions (typically around pH 4) [39]. Unfortunately, mild heat treatment causes protein loss and partial denaturation while the final precipitation step may result in full protein denaturation and hence loss of solubility and functionality. Within the last decade, several improvements to the conventional refinery process have been introduced. This includes addition of e.g. pH stabilizing buffer components [40] or sulphites to reduce oxidative enzymatic activity [41–43]. Moreover, membrane-based approaches are beginning to emerge as a milder and non-destructive biorefinery approach compared to the conventional [42–44].

While membrane-based approaches also have high theoretical potential in green biorefining [45,46], they are typically challenged by especially fouling, limiting protein transmission and thereby reducing efficiency and yields [15,47,48]. Pre-treatments such as mild heat treatment combined with centrifugation have been proposed, albeit at the cost of protein loss and partially impaired functional properties [12,43,47,49]. Recently, a green biorefinery concept based on multi-stage cross-flow membrane filtration was described, where the combination of multiple filtration operations and the use of cross-flow membrane modules under carefully selected parameters mitigates the challenges of membrane fouling to a very large extent [50,51]. This process does not include the use of any heating, centrifugation, or acid/alkaline treatments, thereby facilitating production of a fully native and highly functional protein product [51,52]. However, producing native protein from leaves is still challenged by the very dynamic environment generated immediately after wet fractionation. In the green juice, proteins and phytochemicals that are not co-compartmentalized *in vivo*, become mixed. This facilitates a plethora of potential reactions of both spontaneous and enzymatic nature. For instance, plant proteases obtain access to new protein substrates initiating proteolytic degradation [16,53,54], while plant phytochemicals such as polyphenols may interact with proteins. These interactions may cause modifications that hamper functionality, digestibility, and sensory attributes as well as potentially facilitating protein cross-linking, which may ultimately lead to aggregation, precipitation, and protein loss [55–57]. Controlling these potentially detrimental reactions is key for producing a high-quality native protein product, as proteins and enzymes are not immediately denatured and isolated, but remain functional and active throughout the process [58].

Most research thus far has, with good reason, focused on RuBisCO as the main constituent in protein isolates from green leaves, and very little attention has been given to quantitative composition of the remaining ∼50% of the proteome in green juice [59–61], let alone in downstream refined protein products. On the plant-level, perennial forage crops such as grasses, clovers, and alfalfa, have in contrast been extensively studied for protein-level composition. However, this research has predominantly focused on understanding plant physiology, metabolism, and response to external stress factors [62–69]. Similarly, other leafy greens like sugar beet leaves [70], orange leaves [71], and spinach [72] have been characterized for their protein composition, albeit a protein-level understanding of their processing is lacking. While the high content of RuBisCO makes it a key factor in understanding e.g. bulk functionality of leaf protein as a food ingredient, the remaining proteome, let alone how it is affected by applied processing and refinement methods, will nevertheless have a large impact on the bulk properties.

The aim of this work was to address this knowledge gap by in-depth characterization across unit operations within a membrane-based green biorefinery concept [51]. In the process, a macerated green biomass, here a clover and grass blend, is initially mixed with a stabilizing buffer and wet fractionated in a screw press. The resulting green juice is then refined using multi-stage cross-flow membrane filtration: Firstly, green juice is separated in the first filtration stage wherein the retentate represents a “green protein” suitable for feed. The permeate is further separated in the second filtration stage and subsequent diafiltration to remove small molecule contaminants such as free phytochemicals as well as stabilizers and salts from the added buffer. This produces a concentrated product stream with a high protein content and void of green color and grassy smell and taste [51]. As temperatures are kept relatively low (5-35 °C) throughout the process and pH is maintained at a near-neutral level, proteins are retained in their native form thereby rendering the process non-destructive. This, in turn, ensures a high solubility across a wide pH range as well as retaining the endogenous functionality of the proteins [52]. In this work, mass spectrometry-based bottom-up proteomics was applied to perform an in-depth and quantitative characterization of the proteome across all stages and unit operations within the green biorefinery concept. This approach enables a deeper understanding of the proteome composition and potential, as well as insights into protein-level selectivity throughout the membrane-based processes. The quantitative characterization furthermore enables the coupling between protein-level composition and bulk attributes of the protein extract/isolate. Pigments content was quantified to support interpretation and enhance understanding of the processes. Overall, this study provides a foundation for advancing green biorefining by identifying specific requirements and possibilities for optimization of the concept, ultimately improving yields and quality of the protein products as food ingredients.

## 2. Materials and methods

### 2.1. Biomass production and processing

Clover grass biomass was harvested on August 15^th^ 2022 in Denmark (location: 57°02’16.5”N 9°58’42.4”E) and corresponds to the “W33” experiment previously described [51]. Briefly, the biomass originated from the commercial blend “Seed Mix 35” (DLF Seeds A/S, Denmark) consisting of 13% white clover (*Trifolium repens*) and 87% ryegrass (*Lolium perenne*) of different cultivars, but red clover (*Trifolium pratense*) was also observed in the field. Biomass was harvested using an Agillo PXC Solo bush trimmer (Einhell, Germany) pre-bloom. Harvest was performed in the morning, and the leaves (15 kg crude mass, 2.1 kg DM) were transported for further processing within one hour. The biomass was mixed with a stabilizing solution [50,51] in a 1:2 ratio (w/w) and shredded for a minimum of five minutes using a HMI012 Blender (Hamilton Beach Commercial, USA) at 20°C and maximum power. The suspension was immediately thereafter transferred to a KP6 screw press (Vincent, USA) for wet fractionation at 3.3 rpm and 4 bar counter pressure.

The resulting green juice was then subjected to a multi-stage filtration process, comprising initial removal of fiber and particles by passing the juice through a stainless-steel strainer (1x1 mm square holes) followed by sequential bag filtration using BP-420-x series polypropylene bag filters (Spectrum Filtration, India) with nominal pore sizes of 50 and 1 µm. The permeate was then transferred to the filtration unit equipped with a ceramic HTM membrane (60 nm nominal pore size. LiqTech, Denmark) and filtered by cross-flow filtration using a 3 m/s cross-flow velocity and a quasi-constant trans-membrane pressure (TMP) of 0.3-0.4 bar until a volume concentration factor (VCF) of 4.7 was reached. The permeate was then subjected to a second-stage cross-flow filtration using an ST Sanitary 10 kDa polymeric PES membrane (Synder Filtration, USA) at 2 bar TMP until a VCF of 4.6 was reached. In the same rig, the retentate was washed by continuous diafiltration (DF) using a diafactor (i.e. the volume of wash water to permeate volume) of 9.8. The volumetric cross-flow during the second-stage cross-flow filtration and subsequent DF was maintained at ≈2.9 m^3^/h. For further details on the filtration method and setup, please refer to [50,51].

To characterize process-induced changes of the protein composition, sample aliquots were collected at all possible sampling points during the process and frozen at -18°C until analysis. The sampling points represent:

- The unprocessed (lyophilized and cryo-ground) clover grass (Biomass)
- The green juice immediately after wet fractionation (Raw Juice)
- The green juice (after bag filtering) used as feed for the first-stage membrane filtration (S1 Feed)
- The retentate from the first-stage membrane filtration (S1 Ret)
- The permeate from the first-stage membrane filtration (S1 Perm) - also representing the feed solution for the second-stage membrane filtration
- The retentate from the second-stage membrane filtration (S2 Ret)
- The retentate from the diafiltration operation, representing the final diafiltered concentrate stream (DF C) i.e. the food product stream.
- The permeate from the second-stage membrane filtration (S2 Perm)
- The permeate from the diafiltration operation (DF Perm)

A replicate of the final product stream (DF C) was included as validation of protein-level consistency. This sample was extracted from a different container (but from the same batch) as the DF C sample and labelled as “DF C_2”

### 2.2. Initial characterization of process streams

All streams were initially characterized to determine dry matter, crude protein content, optical characteristics (including pigment analysis), and protein composition analysis by SDS-PAGE.

#### 2.2.1. Dry matter and crude protein content

Stream dry matter content was determined gravimetrically following overnight drying (until completely dry) at 105°C. Dry matter was calculated using the determined wet and dry mass of the streams. DM determinations were performed in at least two replicates. The crude protein (CP) content was estimated based on nitrogen (N) content by elemental analysis. N content was determined using a FlashSmart™ CHNS/O Elemental Analyzer (Thermo Fisher Scientific, Germany) operated in CN mode with helium as carrier gas at a flow rate of 140 mL/min. The combustion temperature was set to 950 °C and the temperature of the detector oven was 50 °C. Acetanilide (OAE Labs, UK) was used for instrument calibration for solid samples while a mixture of urea and glucose was used for liquid samples. CP was calculated using an N-to-CP conversion factor of 6.25 [59] and determined in duplicate.

#### 2.2.2. SDS-PAGE

One-dimensional reducing SDS-PAGE analysis was carried out using SurePAGE 4-20% polyacrylamide gels (Genscript, USA) and a Tris-MOPS SDS Running Buffer system (Genscript), as previously described [73,74]. The unprocessed biomass was cryogenically ground and lyophilized while remaining streams were used in their liquid form. The dry biomass was directly mixed with 4x SDS sample buffer (50 mM Tris pH 6.8, 2% SDS, 10% glycerol, 0.02% bromophenol blue, 12.4 mM EDTA, and 50 mM DTT) to a protein concentration of 4 mg/mL (based on CP), whereof 5 µL was loaded after boiling and centrifugation. For liquid sample, a volume corresponding to 20 µg protein (based on CP) was mixed 1:1 with 4x sample buffer. After boiling, the full volume (avoiding any potentially sedimented solids) was loaded to ensure a theoretically equal protein load. 5 µL PIERCE Unstained Protein MW Marker was used for size estimation and electrophoresis was carried out at 150 V for 40-50 min, staining with Coomassie blue and imaging using a ChemDoc MP imaging System (BioRad, USA).

#### 2.2.3. Optical characterization and pigment quantification

The optical properties of the liquid streams were investigated by UV/Vis spectrophotometry using a DS-11 FX microvolume spectrophotometer (Denovix, USA) measuring the UV/Vis absorbance spectrum from 230-750 nm. Samples were analyzed undiluted as the microvolume analysis is not impaired by the usual restrictions for the Beer-Lambert law in spectrophotometry. Spectra were normalized to their maximum absorbance for comparison. The unprocessed biomass was not characterized as this stream is a dried solid.

Additional UV/Vis analysis was performed for all samples to estimate the content of chlorophyll (a and b) and total carotenoids (xanthophylls and carotenes) by aq. acetone extraction according to [75,76] with minor modifications. For dry (Biomass) samples, 30 mg was directly extracted in 300 µL cold 80% aq. acetone (VWR, Denmark) by grinding in a chilled mortar with addition of 10 mg MgO (Sigma Aldrich, Germany) to prevent pheophytin formation. The slurry was transferred to a centrifuge tube, whereafter the mortar was washed with an additional 100 µL 80% aq. acetone, which was subsequently pooled with the extract. This procedure was repeated twice to minimize losses. For liquid samples, 20 µL was mixed with 180 µL cold 90% aq. acetone, vortexed and incubated on ice for 20 minutes for extraction. Following pigment extraction, all samples were centrifuged at 13.4k rcf for 10 minutes at 4°C to clear residual plant debris. Pigment extracts were analyzed by UV/Vis absorbance measured from 300-750 nm using an SDS-11 FX microvolume spectrophotometer. The conventional absorbance range restriction (0.3-0.85 at 663.2 nm, 646.8 nm, and 470.0 nm [75]) was not considered due to using microvolume analysis. All streams were analyzed in triplicates.

The content of chlorophyll a (eq. 1), chlorophyll b (eq. 2), and carotenoids (eq. 3) was calculated according to [75] as:

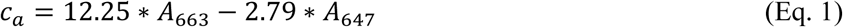

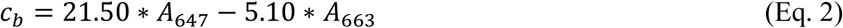

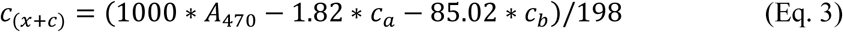

Where c_a_, c_b_, and c_(x+c)_ are the concentrations in the extracts (µg/mL) of chlorophyll a, chlorophyll b and total carotenoids (xanthophylls and carotenes), respectively, and A is the absorbance at the specified wavelength. Pigment content was calculated relative to DM content and CP content of the streams. Due to the low DM and high proportion of non-protein-N in S2 Perm and DF Perm, pigment quantification in these streams was considered unreliable.

### 2.3. Proteomics analysis by bottom-up-proteomics

All streams were subjected to in-depth qualitative and quantitative analysis using mass spectrometry (MS)-based bottom-up proteomics (BUP). Prior to triplicate analysis, the protocol for protein extraction and sample preparation was optimized to the different biomass streams.

#### 2.3.1. Optimization of protein extraction for proteomics analysis

Protein extraction and sample preparation for MS-based BUP analysis was conducted using a combination of the iST kit for plant tissue (PreOmics, Germany) and focused ultrasonication-assisted extraction, as previously described [74]. As representatives of the different streams, the unprocessed biomass, raw green juice, S1 Feed, and S1 Ret were used. All downstream samples were fully soluble in aq. solution and free of particulate matter, why focused ultrasonication was considered unnecessary for these streams. Briefly, an amount/volume of each stream equivalent to 400 µg protein (by N*6.25) was mixed with 600 µL iST “Lyse” buffer and incubated at 95°C and 1000 rpm for 5 min. The suspensions (including solids/suspended matter) were transferred to 1 mL adaptive focused acoustics (AFA) tubes (Covaris) for ultrasonication/AFA-assisted extraction. Prior to the first AFA-assisted extraction cycle (peak incident power of 75 W, a duty factor of 10%, 200 cycles per burst, and 180 s per cycle at 6°C), and following 1, 2, and 4 cycles, 100 µL aliquots were extracted and incubated again at 95°C and 1000 rpm for 5 min. Protein extraction efficiency was evaluated by means of SDS-PAGE analysis (as described above) using ∼13 µg protein loads (assuming full extraction) and by the number of peptide identifications in MS-based BUP analysis, as described below.

#### 2.3.2. Sample preparation and LC-MS/MS analysis of process streams

Based on the optimization experiment, all samples were prepared using the iST kit for plant tissue without AFA-assisted protein extraction. Briefly, a sample mass/volume corresponding to ∼80 µg protein (based on CP) was mixed with Lyse buffer to a final volume of 100 µL. To ensure a correct concentration of Lyse buffer, liquid samples were mixed 1:1 with 2x Lyse buffer and made up to 100 µL by adding 1x Lyse buffer. After protein extraction, reduction and alkylation as described above, the protein extract was mixed with resuspended Trypsin/LysC and digested for three hours on a thermomixer at 37°C and 500 rpm. Subsequently, the digest was transferred to iST cartridges for desalting and cleanup, as previously described [74]. Desalted digest was dried down in a SpeedVac (Thermo-Fisher Scientific, USA) and subsequently resuspended in LC Load and digest concentration was estimated using microvolume UV-spectroscopy (A280) on an SDS-11 FX (Denovix, USA) with standard settings. Samples were diluted as required prior to proteomics analysis. All samples were prepared in triplicates and stored at -18°C until analysis.

LC-MS/MS analysis was performed as shotgun bottom-up proteomics using an EASY nLC-1200 ultra-high-performance liquid chromatography system (Thermo) with ESI coupled to a Q Exactive HF tandem mass spectrometer (Thermo). 1 μg digest was loaded onto a PEPMAP trap column (75 μm x 2 cm, C18, 3 μm, 100 Å) followed by separation on a reversed-phase PEPMAP analytical column (75 μm x 50 cm, C18, 2 μm, 100 Å). Solvent A (0.1 %(V/V) formic acid) and solvent B (80 %(V/V) acetonitrile, 0.1 %(V/V) formic acid), were introduced through a stepwise, 60-minute gradient from 5 % to 100 % solvent B. Data was acquired in full MS/ddMS2 Top20 data-dependent positive mode, employing an MS1 range of 300-1600 m/z. The MS1 and dd-MS2 resolution was set to 60,000 and 15,000, respectively. Automatic Gain Control (AGC) target and maximum injection time were set to 1e5 and 45 ms, respectively, while a 1.2 m/z isolation window and a collisional energy of 28 eV were employed. Dynamic exclusion window was set to 20.0 s, peptide match was set as “preferred”, and “exclude isotopes” was enabled.

#### 2.3.3. Raw data processing

LC-MS/MS data was analyzed in MaxQuant v2.2.0.0 [77,78]. The biomass originates from a field with a mix of perennial ryegrass and white clover, but red clover was also observed. Therefore, all UniProt entries for *L. perenne* (taxid: 4522, 825 entries), *T. repens* (taxid: 3899, 477 entries), and *T. pratense* (UP000236291, 60,146 entries) were used as reference protein databases for protein identification. As *L. perenne* and *T. repens*, the main constituents in the biomass, do not have full proteomes, the reference proteomes for the related *Brachypodium distachyon* (purple false brome, UP000008810, 44,786 entries) and *Trifolium subterraneum* (subterranean clover, UP000242715, 36,725 entries) were used to increase depth of the analysis. Fescues are often found in Danish clover grass fields [79], and as such, all entries from the *Festuca* genus (taxid: 4605, 1,498 entries) were also included. All protein lists were downloaded from Uniprot [80] on December 2^nd^ 2024.

In MaxQuant, oxidation of methionine and protein N-terminal methylation were included as variable modifications while carbamidomethylation of cysteine was defined as a fixed modification. Protein identification was performed using *in silico* tryptic digest, allowing up to two missed cleavages. Peptide size restrictions were defined using a minimal length of seven amino acids and a maximum mass of 4600 Da. On both protein- and peptide-level, quality-based filtering was performed using a 1% false discovery rate. To boost identification rates, matching between runs and dependent peptides were enabled. Protein quantification was done using both label-free quantification (MaxLFQ [81]) and intensity-based absolute quantification (iBAQ [82]). The mass spectrometry proteomics data have been deposited into the ProteomeXchange Consortium via the PRIDE [83] partner repository with the dataset identifier PXD069787 and 10.6019/PXD069787.

### 2.4. Downstream analysis of proteomic data

MaxQuant output data was analyzed with different complementary workflows to obtain both inter- and intra-sample comparative insights. Prior to any downstream analysis, potential contaminant proteins and false positive (reverted sequences) were removed by categorical column filtering. The MaxQuant output data (proteinGroup) including additional downstream processing and appended information can be found in the supplementary information (Table S1). Proteins identified in at least two of three replicates were considered reproducible and were the basis for a qualitative assessment of overlap between streams. Qualitative assessment was visualized in Venn diagrams using Venny 2.1 (https://bioinfogp.cnb.csic.es/tools/venny/index.html).

#### 2.4.1. Comparative analysis with MaxLFQ data

MaxLFQ quantification data was analyzed in Mass Dynamics 2.0 [84]. To handle missing values, the Mass Dynamics built-in “missing not at random (MNAR)” algorithm was used with a mean position factor of 1.8 and a standard deviation factor of 0.3. Pair-wise differential analysis was performed based on sample triplicates and visualized using volcano plots. Proteins were considered significantly differential if the adjusted p-value was < 0.05 and fold change was greater than two (i.e. log2(FC) > 1 or log2(FC) < -1). The built-in Mass Dynamics ANOVA analysis across all stream replicates were used to find globally differential proteins and data was visualized in a heatmap with protein- and replicate-level clustering using a Euclidian cluster distance of seven. Heatmap representations were done both with and without the inclusion of unutilized, low-protein side streams (S2 Perm and DF Perm).

#### 2.4.2. Downstream bioinformatic analysis of MaxLFQ data

Differential proteins from the pair-wise comparison of S1 Ret (484 enriched protein groups) and S1 Perm (353 enriched protein groups) were further analyzed using different bioinformatic pipelines. For this, only the lead protein (i.e. the Uniprot AC# and sequence for the first majority protein) was used to represent each protein group. The subcellular localization of all 837 lead proteins was predicted using DeepLoc v1.0 [85]. For each protein, only the localization category with the highest prediction score was retained. Based on this, the number of proteins associated with each subcellular compartment was quantified per stream. Ten distinct subcellular localizations were included in the analysis: Cell membrane, cytoplasm, endoplasmic reticulum, extracellular, golgi apparatus, lysosome/vacuole, mitochondrion, nucleus, peroxisome, and plastid. The proportion of enriched proteins in the two streams ascribed to each compartment (% of enriched proteins in the respective stream) was then computed to explore potential underlying subcellular descriptors of protein selectivity during the first filtration stage. Similarly, the proportion (% of enriched proteins) of probable membrane-associated proteins were predicted using TOPCONS2 [86]. A protein was considered membrane-associated if it contained at least one transmembrane (TM) domain without discriminating between cell and compartmental membranes. The term “membrane-associated” will from here on refer to proteins associated to biological membranes by predicted transmembrane regions and does not refer in any way to the membranes used for filtration.

#### 2.4.3. Comparative analysis by intra-sample relative quantification with riBAQ and protein grouping

Within each sample replicate, the intensity-base absolute quantification (iBAQ) intensity was used to estimate relative molar protein distribution. Relative molar abundance (relative iBAQ, riBAQ) was obtained by dividing the protein-level iBAQ with the sum of iBAQs within each sample replicate (in %), as previously described [73,74]. To delimit the dataset to only abundant proteins, filtering was applied requiring reproducible identification (i.e. identified in at least two of three replicates) and an abundance threshold (mean replicate riBAQ > 0.5%) in at least one stream. This reduced the number of proteins from initially 3580 to 61 high abundance proteins. Based on prominent and reoccurring proteins within this subset, the full dataset was revisited to determine the sum of riBAQs for all isoforms of particular proteins or protein families to make more generalizable analysis based on the protein type/family rather than on the individual isoform-level. These groups consist of RuBisCO (rbc), Chlorophyll a-b binding protein (CBP), Photosystem I & II proteins (PI/II), Oxygen evolving system proteins of photosystem II (PII-OES), Non-specific lipid transfer protein (nsLTP), thioredoxins (TRX), superoxide dismutase (SOD), ferredoxin (FNR), histones (His), and ATP synthases (ATPs). Lists of lead protein accession numbers (AC#s) for protein groups included in each type/family group can be found in the supplementary information (Table S2).

Using both individually identified protein groups within the abundant subset and the defined groups, a fold change (FC) analysis was performed based on riBAQ abundance. This analysis was performed to investigate protein-level and group-level enrichments across the two filtration stages. For the first filtration stage, the FC was computed by pair-wise comparing S1 Feed, S1 Ret, and S1 Perm. For the second filtration and DF stages, the FC was calculated by pair-wise comparing S1 Perm, S2 Ret, and DF C. Permeate streams from S2/DF were not considered due to their low protein content and hence, unreliable quantification. FC between two streams (A and B) was determined as the log2 transform of the ratio between riBAQs (B/A). Negative values represent enrichment in stream A, while positive values represent enrichment in stream B. Based on the log2 transformation FC < -1 and FC > 1 represent a relative (i.e. part of the whole) two-fold enrichment in the respective stream and a 50% reduction in the other. For proteins not detected in a particular stream, log2FC was set to +/- 10 based on maximal values observed in the analysis. If a protein was not found or had a riBAQ < 0.01% in both streams, log2FC was set to zero.

#### 2.4.4. Bioinformatic analysis of riBAQ data

Different bioinformatic pipelines were developed to obtain further qualitative and quantitative insights on the protein composition in the different streams. All analyses were performed using Python (v3.9.18), supported by a range of open-source libraries: Scikit-learn (v1.3.0) [87] for machine learning, Biopython (v1.78) [88] for biological sequence analysis, Pandas (v2.0.3) [89] for data manipulation, Matplotlib (v3.7.2) [90] and Seaborn (v0.12.2) [91] for visualization, SciPy (v1.10.1) [92] for statistical computations, and NumPy (v1.24.3) [93] for numerical operations.

The dataset initially contained 3,580 protein identifications. A preliminary filtering step removed 70 entries that included non-standard amino acids, resulting in a final dataset of 3,510 protein groups. For groups containing multiple protein IDs, only the first ID was retained to represent that unique identification. Protein sequences corresponding to these IDs were retrieved from the respective FASTA files. For each stream, riBAQ values across three replicates were averaged to compute the mean riBAQ, which was subsequently used to determine protein presence per stream.

##### 2.4.4.1 Analysis of physicochemical properties

A range of sequence-derived features was computed using the *ProteinAnalysis* tool from Biopython (v1.78). These included molecular weight, aromaticity [94], instability index [95], GRAVY score [96], and isoelectric point. Additionally, the predicted fractions of amino acids forming secondary structure elements (helices, turns, and sheets) were calculated. For clarity, all these descriptors, whether purely chemical or structural predictions, are collectively referred to as physicochemical properties, or simply “properties” in this work.

##### 2.4.4.2 Unweighted analysis

Each protein was treated as an independent data point and the distribution of physicochemical properties across streams was initially explored using boxplots. To evaluate if there were significant differences between streams properties, a one-way ANOVA was performed using SciPy (v1.10.1). Subsequently, dimensionality reduction was applied using Truncated Singular Value Decomposition (*TruncatedSVD*) from scikit-learn (v1.3.0), allowing the data to be represented in a two-dimensional space for visualization. Clustering analysis was then performed on the normalized properties. The optimal number of clusters was estimated using the elbow method and the Calinski–Harabasz index. Clusters were defined using the K-Means algorithm. The 2D representation of the data was then visualized to highlight groupings based on clusters, stream assignments, and predicted subcellular localizations.

##### 2.4.4.3 Abundance-weighted analysis

In this analysis, proteins identified in each stream and replicate were considered as independent observations. To incorporate protein abundance, the physicochemical properties of each protein were multiplied by its corresponding riBAQ value. The resulting weighted values were then summed across all proteins within each stream and replicate, yielding 30 aggregated data points (10 streams × 3 replicates), each represented by a vector of weighted physicochemical properties. As in the unweighted analysis, the distribution of these weighted properties across streams was visualized using boxplots and statistically evaluated using one-way ANOVA. Dimensionality reduction was applied to the normalized weighted data, followed by clustering analysis. The results were projected into a two-dimensional space for visualization. Additionally, the distribution of physicochemical properties within each cluster was examined using boxplots to reveal cluster-specific trends and potential biological/process-induced relevance.

##### 2.4.4.4 Analysis of subcellular localization and transmembrane regions

Protein subcellular localization was predicted for each experimental stream, and for all 3,510 proteins using DeepLoc v1.0 [85]. A protein was considered present in a specific stream if it was identified in at least one of the three replicates. Based on this, the number of proteins associated with each subcellular compartment was quantified per stream. This yielded a matrix representing 10 streams, each characterized by the distribution of proteins across the ten compartments. To explore underlying patterns, dimensionality reduction was applied to visualize the data in two dimensions. Clustering was subsequently performed on the complete dataset, and the resulting groups were visualized within the same 2D space to facilitate interpretation of stream-specific localization trends. Probable membrane-associated proteins were predicted across all identified lead proteins (3510 after filtering) using TOPCONS2 [86], as described above. The abundance of protein-based subcellular localization and membrane association was subsequently computed using riBAQ values within each stream.

### 2.5. Statistical analysis

Data from dry matter, crude protein, pigment analysis and number of protein identifications was analyzed to identify statistical differences using GraphPad Prism (v.10.0.2, build 232). Comparison of means from the different streams was performed as ordinary one-way ANOVA and Tukey’s multiple comparison testing using a 95% confidence interval and assuming Gaussian distribution of residuals and equal standard deviations between groups of unmatched replicates. All p-values are reported as adjusted p-values to account for multiple comparisons.

## 3. Results and discussion

During the entire process, a variety of streams were sampled for investigation. Initially, dry matter (DM) and crude protein (CP) content was determined (Table 1) as the basis for additional analysis and to investigate the fate of the bulk protein and dry matter over the process.

**Table 1:** Overview of sampled streams and their dry matter (DM) and crude protein (CP) content. For each stream, the short name and stream description are indicated along with DM (%) and CP (% of DM) using a N-to-protein conversion factor of 6.25. Values are given as means ± standard deviation (n = 2). Different lower-case superscript letters (within each column) represent significant differences between means based on one-way ANOVA and Tukey’s multiple comparison testing with 95 % confidence intervals (p < 0.05). For additional results and mass balances for DM and CP over the entire season, see Matsson *et. al* (2025).

| Stream | Description | Dry matter (%) | Crude protein (%DM) |
| --- | --- | --- | --- |
| **Biomass** | Unprocessed biomass from harvest | 14.2 ± 2.15 <sup>a</sup> | 25.0 ± 0.74 <sup>c</sup> |
| **Raw Juice** | Green juice after wet fractionation | 3.42 ± 0.02 <sup>c,d</sup> | 20.6 ± 0.36 <sup>d</sup> |
| **S1 Feed** | Green juice after bag filtering (feed for first filtration stage) | 3.36 ± 0.02 <sup>c,d</sup> | 20.4 ± 0.05 <sup>d</sup> |
| **S1 Ret** | Retentate from first filtration stage (animal feed protein stream) | 5.48 ± 0.01 <sup>b</sup> | 38.7 ± 0.67 <sup>b</sup> |
| **S1 Perm** | Permeate from first filtration stage (feed for second filtration stage) | 2.37 ± 0.35 <sup>d,e</sup> | 10.3 ± 0.20 <sup>e</sup> |
| **S2 Ret** | Retentate from second filtration stage ("feed" for diafiltration) | 2.14 ± 0.00 <sup>e</sup> | 21.1 ± 0.87 <sup>d</sup> |
| **S2 Perm <sup>1</sup>** | Permeate from second filtration stage (side stream) | N/A | N/A |
| **DF Perm <sup>1</sup>** | Permeate from diafiltration (additional side stream) | N/A | N/A |
| **DF C** | Concentrate from diafiltration (final food protein stream) | 0.48 ± 0.00 <sup>f</sup> | 57.7 ± 0.86 <sup>a</sup> |

<sup>1</sup>Dry matter and crude protein was not recorded for the permeate streams from the second filtration and diafiltration stages.

A substantial and highly significant (p < 0.0001) loss of DM is observed in all streams compared to the unprocessed biomass (Table 1, Fig. S1A). Moreover, a significant (p < 0.0032) accumulation of DM in the S1 retentate is observed compared to both upstream (Raw Juice and S1 Feed) and downstream sampling points. This can be attributed to retention of particulate matter and aggregates permeating the initial police filtering stage between Raw Juice and S1 Feed, where no significant (p > 0.99) DM loss was observed. While a notable DM loss was observed between S1 Feed and S1 Perm due to the retention of larger components in S1 Ret, the difference was not significant (p = 0.11), likely due to the low number of sample replicates (n = 2). Nevertheless, compared to the S1 Feed, additional and significant decreases of DM content were observed moving further downstream to the S2 Ret (p = 0.047) and finally the DF C stream (p = 0.004). These findings are in good agreement with the general trends observed across a full season of membrane-based biorefining [51]. This process produces a native and fully soluble product with CP ranging from 57% to 73% (DM basis). While the experiment presented here represents a lower purity concentrate compared to other runs within the same season, the protein composition (Fig. 1A) is comparable to concentrates obtained under similar conditions based on previous SDS-PAGE analysis of the final DF C stream [51].

**Figure 1:**
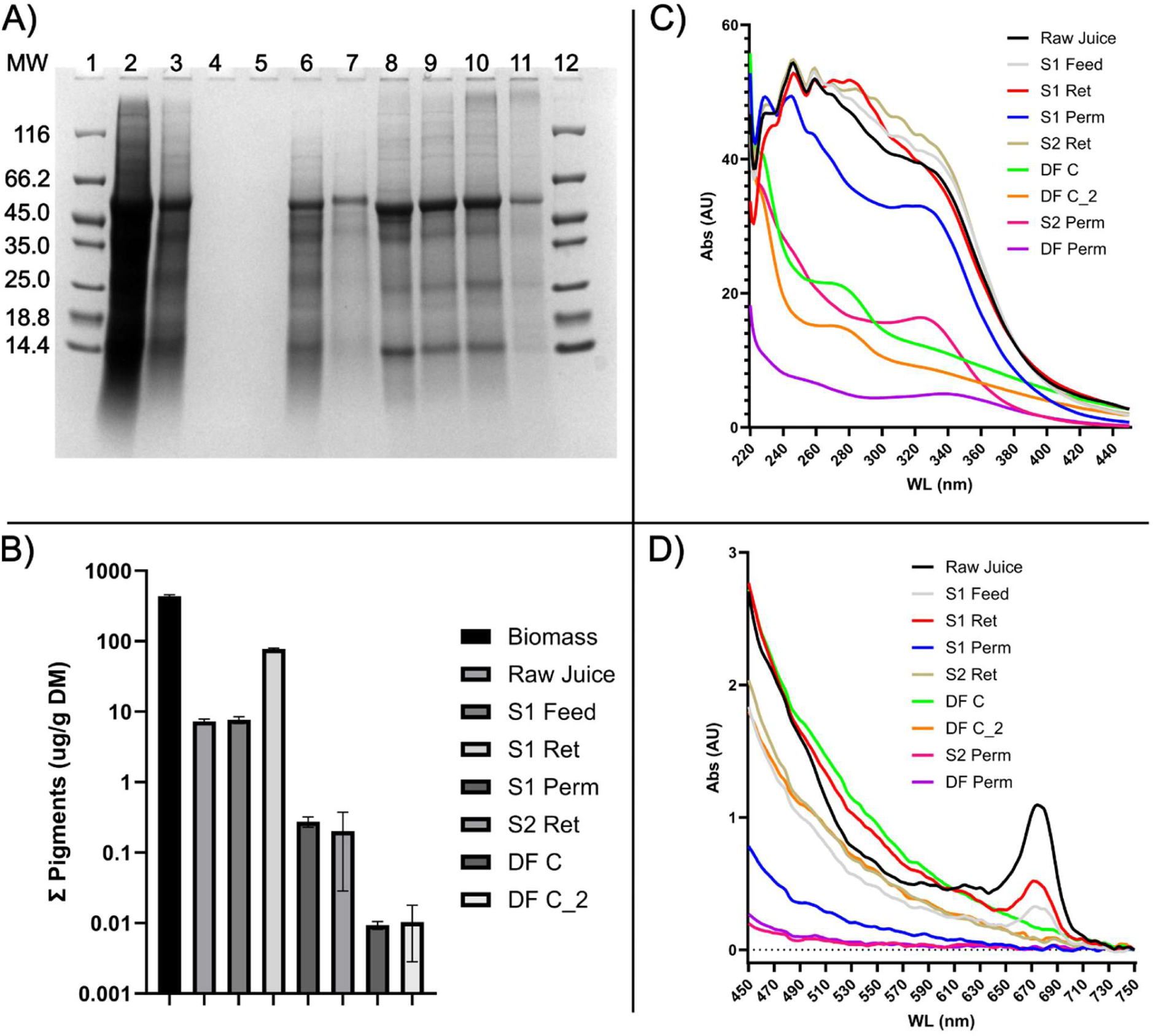
Crude protein and pigment/color characterization. A) Reducing SDS-PAGE analysis of the different streams represented by unprocessed biomass (2), Raw juice (3), S1 Feed (4), S1 Ret (5), S1 Perm (6), S2 Ret (7), S2 Perm (8), DF Perm (9), DF C (10), DF C_2 (11). MW marker is loaded in wells 1 and 12 and protein MW (kDa) annotated. B) Quantification of pigment content in the different stream by UV/Vis following acetone extraction. For each stream, the sum of pigments (chlorophyll a+b and carotenoids (xanthophylls and carotenes)) relative to the stream dry matter content is shown using a Log10 scale to accommodate the large dynamic range of values. Quantification of individual pigments is presented in supplementary materials (Fig. S2). C) Absorbance spectra of liquid streams in the range 220-450 nm. D) Absorbance spectra of liquid streams in the range 450-750nm. Absorbance spectra (C-D) are presented in two separate plots due to the high UV absorbance in the streams.

While the DF C stream has a significantly higher CP content (and thereby higher protein purity) compared to all other streams (Table 1, Fig. S1B), it also contains lower DM compared to all other upstream sampling points, which can be attributed mainly to two aspects. Firstly, the biorefinery concept uses a stabilizing buffer (1.7% DM) to ensure a more consistent process over time. The stabilizing components, such as buffer salts, are washed out during the second filtration and subsequent diafiltration (DF) stages, thereby directly reducing DM [51]. Secondly, a substantial loss of protein can be observed over each of the unit operations starting already from the wet fractionation, where approximately 60% of the protein (by CP) was retained in the fiber fraction (pulp) [51]. Moreover, a substantial loss of nitrogen (approximately 25%) was observed in the downstream permeates (S2 Perm and DF Perm). This is ascribed to loss of very small proteins, peptides and partially degraded protein, and particularly non-protein nitrogen of both organic and inorganic origin. Ultimately around 7% of the initial protein (by CP) end up in the final product stream. This yield was in fact the lowest of the production season, where CP yields in DF C range from approximately 7% to 15% [51], but despite a low yield and purity, this specific batch was still chosen for in-depth characterization based on the representative parameters used during the membrane-based biorefining process and the availability of subsamples from all process streams. While other methods for estimation of protein content in the streams could have been explored, previous studies have shown these to be incompatible with the workflow [59]. For instance, optical methods such as BCA, A280, and the fluorescence-based Qubit assay are impaired by the high level of interference from pigments and other UV-active species present in many of the analyzed streams, leading to inaccurate protein quantification [73,97,98]. Their successful application would require initial and selective protein isolation to remove interferences, which may introduce additional bias and/or result in partial protein denaturation and precipitation. As such, only N-based protein estimation was used in this study. While using the Jones N-to-protein conversion factor of 6.25 [99] has been deemed inaccurate in plant samples including grass species [100–102], stream-specific factors have not yet been determined. As such, using the Jones factor was employed for methodological consistency.

### 3.1 Efficient pigment removal and apparent changes to protein composition

While the overall protein profile appeared largely conserved along the process, based on SDS-PAGE analysis (Fig. 1A), distinct changes were observed. The profile of the two upstream samples (Raw juice and S1 Feed) as well as the S1 Ret resembled that of the unprocessed biomass (lanes 2-5). Downstream of the first filtration stage in the S1 Perm (lane 6), high MW proteins (> ∼100 kDa) appeared largely depleted. Several bands become visible again in downstream samples, potentially due differences in salt concentrations. In contrast, apparent high-MW aggregates, observed at the top of the lanes in upstream samples, appeared selectively depleted. This observation underlines the size-based selectivity of the first stage filtration. Interestingly, the distinct band at ∼25 kDa seen in upstream samples appears to be retained in the first stage filtration, as this is absent in all samples downstream (lanes 5-11). This indicates additional selectivity in the first filtration operation that cannot be simplified to monomeric protein MW. This observation is in agreement with an earlier study, where removal of this particular band was also observed [51]. The removal of a 25 kDa protein using a 60 nm membrane in the first stage filtration indicates that this particular protein is likely found within larger supramolecular complexes or in aggregated form in the green juice. Overall, the protein profile in the streams resembles that obtained in previous studies on green biomass from not only grass but also other leafy greens [52,103–105]. The different appearance of the two DF C samples can be attributed to differences in protein load.

During SDS-PAGE analysis, a green hue was also observed below lanes containing samples upstream from the first filtration stage and in S1 Ret. While this observation is not reflected post imaging of Coomassie-stained gels, it agrees with a distinct color change in the liquid streams, likely ascribed to chlorophyll depletion. To further investigate this phenomenon, we used UV/Vis analysis of the streams to both characterize their optical properties as well as to specifically quantify pigments. For the three quantified pigments (chlorophyll a (chl a), chlorophyll b (chl b), and total carotenoids (cart) (i.e. xanthophylls and carotenes), similar trends were observed (Fig. S2 A-C), also reflected in their sum (Fig. 1B). The unprocessed biomass was found to have relatively high content of chl a (140 µg/g DM), chl b (230 µg/g DM), and cart (68 µg/g DM) for a total pigment content of 435 µg/g DM. This also corresponds to a total chlorophyll content of 370 µg/g DM or 2.6 mg/g fresh biomass (FM). This value is in agreement with previous studies with total chlorophyll typically reported to be 2-4 mg/g FM [106,107] in cultivated ryegrass. Further comparisons with literature are complicated by differences in analytical methods, as many apply SPAD-meters and report in SPAD units [103,108]. The green juice (Raw Juice) has a significantly lower (p < 0.0001) content of all individual pigments and their sum, albeit still at a total level (7.7 µg/g DM), sufficiently high to produce an intense green color of the juice.

The incomplete pigment extraction during wet fractionation by screw pressing agrees with earlier studies, where pigments could be extracted continuously through multiple passes of alfalfa biomass in a screw press [103]. The police filter applied prior to first stage filtration had no significant (p > 0.9999) effect on pigment levels. Over the first filtration stage, a 10-fold enrichment (p < 0.0001) of pigment was observed in S1 Ret (77 µg/g DM) while a 96% depletion (p < 0.0001) was found in the S1 Perm (0.27 µg/g DM). This shows a highly efficient pigment retention in the first filtration stage. Pigment levels were further (27%) reduced during the second filtration stage (S2 Ret: 0.20 µg/g DM) albeit not with statistical significance. However, pigments were continuously removed during DF (DF C: 0.009 µg/g DM), representing a significant (p = 0.046) pigment depletion of 55% from the S1 Perm stream. Overall, this illustrates the highly efficient removal of 99.9% of chlorophylls and carotenoids from the green juice (S1 Feed) to the final product stream (DF C). These trends were also observed when pigments were quantified relative to CP (Fig. S2 D-F).

The efficient pigment removal in the first filtration stage is also directly reflected in the UV/Vis spectra of the streams (Fig. 1C-D). It is noteworthy that a distinct peak at ∼670nm is evident in upstream samples and in S1 Ret (Fig. 1D). This corresponds well with the known absorbance of chl a in this range [108]. A distinct peak for chl b and cart was not evident due to the generally high sample absorbance, which increased towards lower wavelengths. The chl a peak is absent downstream of the first filtration stage, further highlighting the efficient removal in this unit operation. In addition to the phytopigments, the UV absorbance (Fig. 1C) is also very high in upstream samples, indicating high content of UV-active and aromatic compounds. Interestingly, the overall UV absorbance also generally appears to decrease during DF, despite an increasing proportion of CP in downstream samples. More specifically, compounds responsible for absorbance around 230-255nm and 310-350nm are washed out initially in S2 Perm and subsequently in DF Perm. Absorbance at 250nm has previously been ascribed to e.g. flavonoids like quercetin and rutin while absorbance at 325nm has been ascribed to e.g. the phenolic acid, chlorogenic acid [109]. In contrast, the emergence of a peak around 280nm, indicative of protein UV absorbance, can be seen in the final product streams (DF C and DF C_2). This coincides with the substantial concentration of CP during the second filtration and DF stages from 10% in S1 Perm, over 21% in S2 Ret to 58% in DF C. This further substantiates protein enrichment in the product stream in addition to removal of other and undesired UV-active species such as quercetin and phenolic acids, which are associated with lower sensory quality such as bitterness and astringency [30–32]. However, this warrants further investigation.

### 3.2 Optimization of sample preparation for bottom-up proteomics

Prior to in-depth analysis of the different process streams, the protein extraction protocol was optimized by combining focused ultrasonication (AFA) with the PreOmics iST plant tissue kit. To evaluate the effect of ultrasonication, a varied number of AFA extraction cycles (0, 1, 2, 4) was applied combined with the thermochemical extraction from the kit. For this purpose, four different upstream samples (Biomass, Raw Juice, S1 Feed, and S1 Ret) were selected while downstream samples were considered fully soluble. By SDS-PAGE analysis (Fig.S3A+B), there appeared to be no substantial impact of applying AFA ultrasonication as part of the protein extraction workflow regardless of the stream and the amount of solids and residual plant material. In fact, by considering the number of identified peptides as a function of AFA cycles (Fig. S3C), a general decreasing trend was observed with an increasing number of cycles regardless of considering peptide IDs by MS/MS only or by including matching between runs. While the origin of this effect was not investigated further, it may reflect partial protein damage (e.g. by degradation or denaturation-induced aggregation and precipitation) as previously reported [110–113]. These data suggested that the use of AFA did not appear to have a positive effect on this biomass or resulting streams, underlining the importance of optimizing the workflow every time a new biomass is investigated [74]. While earlier studies on green biomass have indeed included AFA as part of the protein extraction workflow [59], AFA was not included in the protein extraction for the full characterization of process streams in this work.

### 3.3 Process-induced qualitative changes in protein composition

Across all samples, a total of 3580 protein groups, representing 8766 potential proteins, were identified after filtering contaminants and false positives. The number of protein IDs is substantially (∼3 fold) higher than in a recent study on biomass and green juice from lab-cultivated *L. perenne* [59]. This increase can be attributed to the biomass in this work originating from a field cultivated with a mixture of perennial grass and clovers. With the exception of the two “side streams” (S2 perm and DF perm), >2000 protein groups were identified in all streams (Fig. 2A). The lower number of protein IDs in the S2 and DF permeates was expected as these represent what is transmitted through the second-stage filtration using a 10 kDa membrane. Moreover, a lower number of protein/peptide IDs in the DF permeate is expected as this represents continued washing after second stage filtration. The fact that proteins were identified in these streams can be ascribed to primarily partially degraded protein (i.e. peptides), as no visible protein was observed by SDS-PAGE (Fig. 1A). Interestingly, a distinct drop and significant (p<0.0001) decrease in number of protein IDs across the first-stage filtration was found. This observation is consistent regardless of considering individual sample replicates (Fig. 2A) or the number of reproducibly quantified (i.e. found in at least two of three replicates) proteins (Fig. S4A). In all samples upstream of the first stage filtration, >2200 proteins were identified (up to 2575 reproducible IDs in the S1 Feed) while the number of protein IDs drops to ∼2000 in S1 Perm. This indicated selectivity in the membrane operation. Interestingly, a downstream increase in protein IDs was observed that continued to increase further downstream in the purification and concentration process. Compared to the S1 Perm (2004 reproducible IDs), the increase in protein IDs went from being insignificant (p=0.337) in S2 Ret (2127) to being significant (p<0.005) in the two replicates of DF C (2232 and 2202). This increase could be ascribed to an increasing protein purity, thereby minimizing interference and noise.

**Figure 2:**
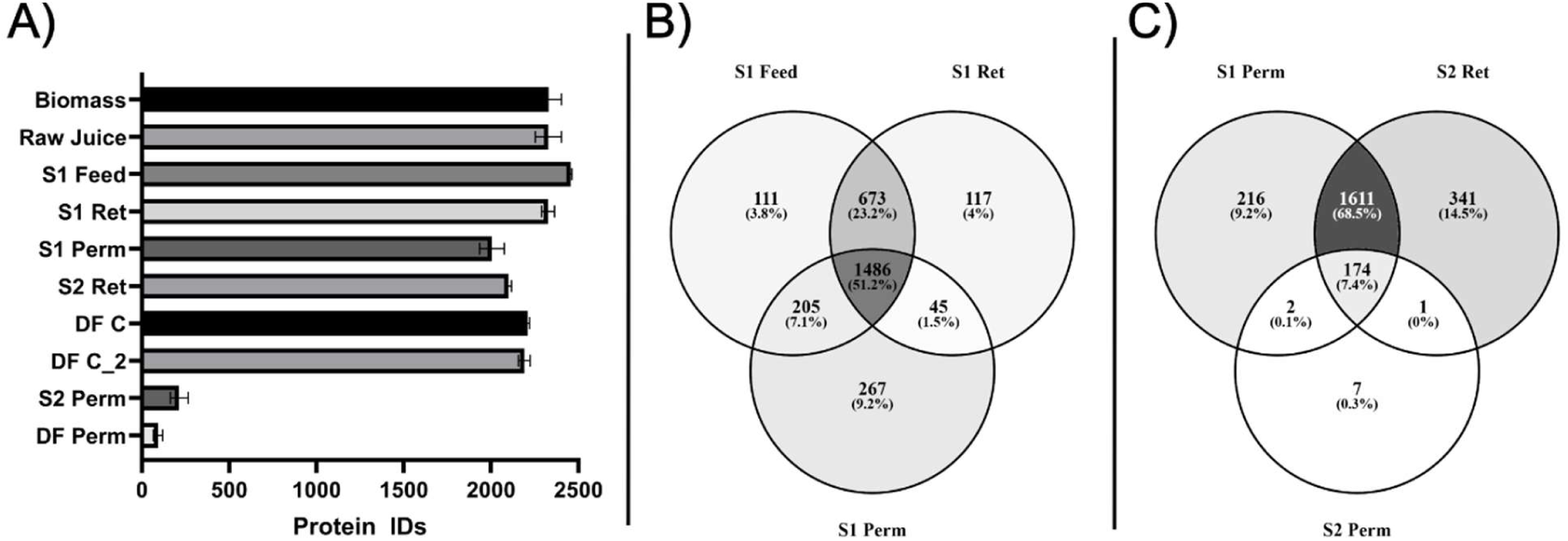
Overview of protein identification and overlap between streams. A) Number of protein IDs for each stream depicted as mean +/- standard deviation from sample triplicates. B) Overlap of reproducibly identified (i.e. in at least two of three replicates) proteins between S1 Feed, S1 Ret, and S1 Perm. C) Overlap of reproducibly identified proteins between S1 Perm, S2 Ret, and S2 Perm. In the Venn diagrams (B-C), both the number of reproducible protein IDs and proportions are shown.

The selectivity across the first stage filtration was also reflected in qualitative data when inspecting the overlap between reproducibly (i.e. in at least two of three replicates) identified proteins for the S1 Feed, S1 Ret, and S1 Perm (Fig. 2B). From the ∼2900 protein IDs found in any of these streams, only 51% (1486) were found in all three streams. 27% (790) of the protein IDs were found in S1Feed and S1 Ret or only exclusive in S1 Ret, indicating that a substantial number of proteins did not appear to cross the first membrane. Furthermore, 16% (472) were found either in both S1 Feed and S1 perm or exclusively in S1 Perm, further substantiating selective transmittance across the first membrane. In comparison, the second stage filtration (Fig. 1C) displayed almost full protein retention by the second stage membrane based on the number of protein IDs. From the 2352 protein identified across the three streams, less than 0.5% (9) proteins were found in the intersect between S1 Perm and S2 Perm or exclusively in S2 Perm. The identification of exclusive proteins in this intersect or in S2 Perm can be ascribed to instrument sensitivity, as all these proteins were of low abundance. In a higher complexity mixture of proteins (i.e. feed side of the second stage membrane), low abundance proteins (i.e. their tryptic peptides) may not be detected due to factors such as e.g. ionization competition, co-elution, and ion suppression [114,115], but that does not mean they are not present on the feed side and suddenly appear on the permeate side. 174 proteins (7%) were found in all three streams indicating partial transmission – or potentially partial degradation and subsequent transmission of peptide products. The remaining proteins were identified in either S1 Perm, S2 Ret or, representing the majority of protein IDs (69%), both.

### 3.4 Distinct and quantitative process-induced enrichment and depletion on the protein level

Using label-free quantification (MaxLFQ), the quantitative proteome changes across the different stages of the membrane-based process were investigated. Based on normalized MaxLFQ data, a clear differentiation in the quantitative protein composition was evident, when considering differentially abundant proteins by ANOVA (Fig. 3A). This was further reflected in the replicate-level Pearson correlation (Fig S4B) and in the principal component analysis (Fig. S4C). Upstream samples (Biomass, Raw Juice, S1 Feed, and S1 Ret) all display quite comparable qualitative and quantitative composition also resulting in a clear sample clustering in PCA (Fig. S4C). Similarly, downstream samples (S1 Perm, S2 Ret, DF C, and DF C_2) are highly comparable and cluster while the residual side streams from the second filtration and DF stages (S2 Perm and DF Perm) represent a third and distinct cluster. The similarity within the clusters is also noticeable when performing hierarchical clustering on the replicate level. When including the two residual side streams (Fig. S4D), clustering of S2 Perm and DF Perm replicates overlap. When omitting these streams (Fig. 3A), DF C and DF C_2 co-cluster. As these are technical replicates but represent the same product batch, this also underlines batch consistency. Similarly, Raw Juice and S1 Feed co-cluster, indicating that the pre-filtering step had very little impact on the quantitative protein composition. Nevertheless, two very distinct clusters are observed at the protein level: One group of proteins enriched in upstream samples and one group enriched in downstream samples (Fig. 3A). This further substantiates the finding by analysis of protein IDs alone, that particularly the first membrane filtration represents a highly selective process. This is also reflected in the optical properties of the samples (Fig. 1) and is in agreement with observations during processing across the whole season [51].

**Figure 3:**
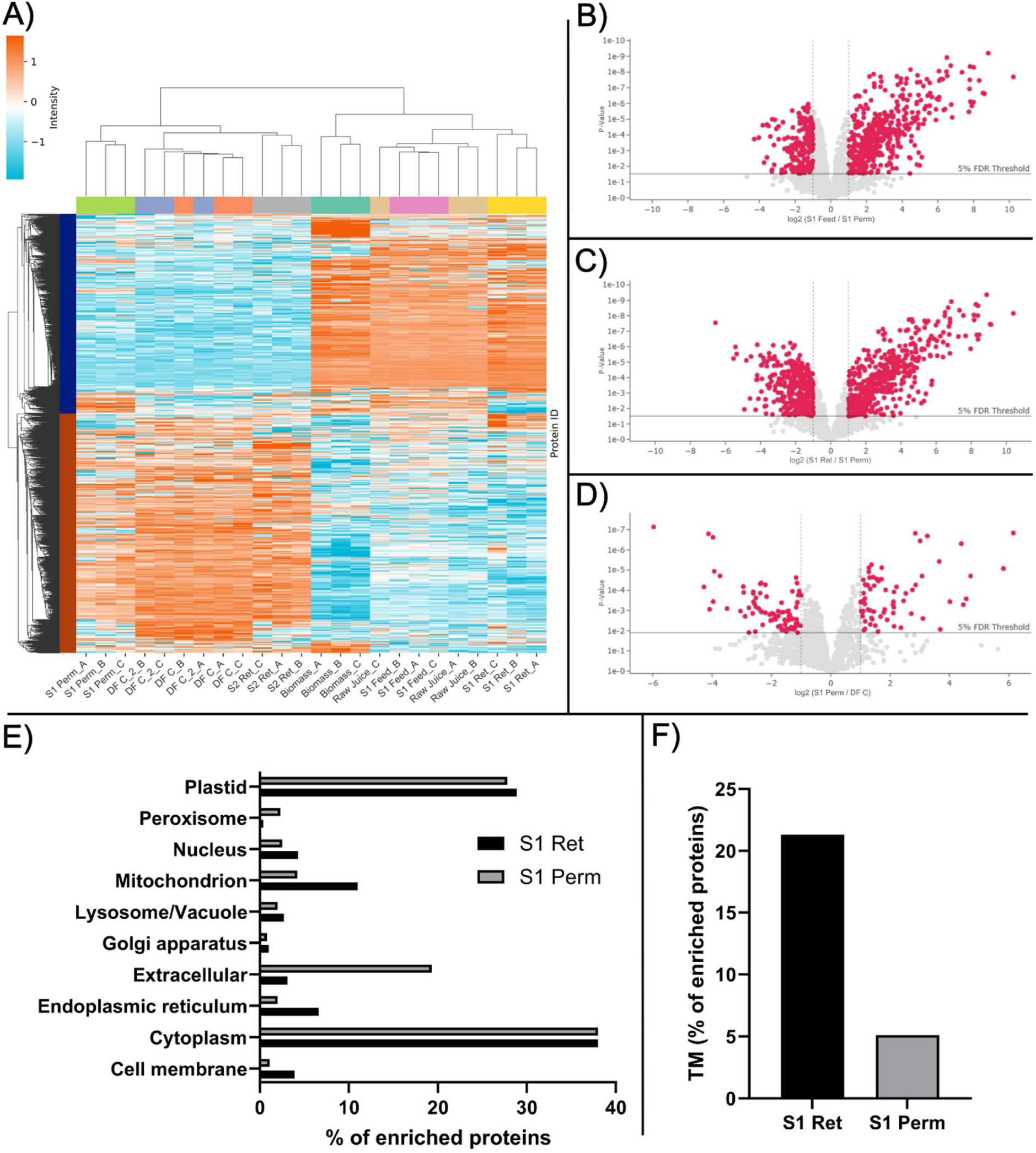
Comparative analysis across streams using label-free quantification (MaxLFQ). A) Heatmap representation of differentially abundant proteins by ANOVA analysis in MassDynamics. Data is depicted as z-score standardized MaxLFQ intensities by row (protein group) and clustered using a Euclidian distance of 7. B) Volcano-plot from pairwise differential analysis of protein LFQ intensities for S1 Feed vs. S1 Perm. C) Volcano-plot from pairwise differential analysis of protein LFQ intensities for S1 Ret vs. S1 Perm. D) Volcano-plot from pairwise differential analysis of protein LFQ intensities for S1 Perm vs. DF C. In all volcano plots Highlighted proteins (in red) were found to be significant (p < 0.05) and substantially (FC ≥ 2) differentially abundant in the pairwise analysis in Mass Dynamics. E) Subcellular distribution of enriched proteins identified from pair-wise differential analysis of S1 Ret vs. S1 Perm. Distribution is depicted as proportion of proteins ascribed to a particular subcellular compartment (as predicted by DeepLoc) relative to all proteins enriched in the particular stream (in %) F) Distribution of enriched membrane-associated proteins (TM) identified from pair-wise differential analysis of S1 Ret vs. S1 Perm. Distribution is depicted as proportion of proteins with transmembrane regions (as predicted by TOPCONS2) relative to all proteins enriched in the particular stream (in %)

To obtain further insight into the selectivity in this step, pair-wise analysis of S1 Feed and S1 Perm as well as S1 Ret and S1 Perm was performed. In the comparison of S1 Feed and S1 Perm (Fig. 3B), 190 proteins were found significantly (adjusted p-value < 0.05) and substantially (fold change ≥ 2) enriched in the permeate while 454 proteins were depleted. Comparing S1 Ret and S1 Perm (Fig. 3C), 353 proteins were enriched in the permeate while 484 proteins were depleted. This corroborates that the biggest differences originating from protein-level selectivity in the first stage filtration comes from comparative analysis of the two products more so than by comparison with the feed. A substantial overlap between enriched or depleted proteins in the S1 Feed vs. S1 Perm comparison were also found enriched or depleted, respectively, when comparing S1 Ret and S1 Perm (Fig. S5), further illustrating the overall comparability of S1 Feed and S1 Ret. Ultimately, lists of the 837 differential proteins from the S1 Ret and S1 Perm pair-wise comparison were exported for further analysis (Table S3).

Subcellular localization prediction with DeepLoc [116] was used to investigate if the enriched proteins in the respective streams were biased toward certain subcellular compartments (Fig. 3E). In S1 Perm, a larger proportion of extracellular (6.2-fold) and peroxisome (5.5-fold) proteins were enriched. However, the low number of enriched peroxisome proteins (10 of 834) makes this compartment associated with larger uncertainty in terms of relative difference in enrichment. In contrast, S1 Ret had a higher proportion of proteins localized in the cell membrane (3.5-fold), endoplasmatic reticulum (3.3-fold), and mitochondria (2.6-fold). Plastid and cytoplasmic proteins showed less than 1.05-fold difference, while modest enrichment (1.2-1.7-fold) in S1 Ret was seen for remaining compartments. These observations indicate that particularly extracellular, and thereby presumably soluble proteins, are more easily transmitted through the first stage filtration membrane, while e.g. cell membrane proteins are retained, as these are potentially still bound to cell membrane fragments or prone to aggregation. To test this hypothesis further, we predicted transmembrane regions for all proteins, as some proteins localized to highly represented compartments such as cytoplasm and plastid may still be membrane-associated. Based on prediction of all membrane-associated proteins, it was evident that proteins with predicted transmembrane regions were 4.2-fold more enriched in S1 Ret (21.3%) compared to those enriched in S1 Perm (5.1%), substantiating a high degree of selective retention of membrane-associated proteins in the first stage filtration (Fig. 3F).

The most highly enriched in S1 Perm, and therefore also likely enriched in the final product, were Nodulin-related protein 1 (A0A2K3JMR2, log2FC=6.6), Non-specific lipid-transfer protein (A0A2K3NE56, log2FC=5.5), and Reticuline oxidase-like protein (A0A2K3N926, log2FC=5.4); all from *T. pratense*. Nodulins are proteins upregulated in legumes because of the symbiotic nitrogen fixation process with *Rhizobium* bacteria [117] but there is no further knowledge on functionality in a food context or safety issues related to e.g. allergenicity of nodulins. Non-specific lipid-transfer proteins (nsLTPs), in contrast, are well-known food allergens with sensitization capabilities. Reticuline oxidase belongs to the class of berberine bridge enzymes (BBE). BBE-like enzymes have also previously been reported to cause allergenic responses in people with pollen allergies [118]. The presence of such proteins in a food product may entail certain precautions in terms of proper labelling.

In S1 Ret, a very different subset of proteins was found highly enriched. The most enriched protein were Chlorophyll a-b binding protein from *T. subterraneum* (A0A2Z6NV37, log2FC=10.4), Photosystem II protein D1 from *L. perenne* (Q95GL2, log2FC=9.1), Photosystem II CP47 reaction center protein from *T. repens* (A0A023HPN2, log2FC=9.1), and Photosystem II D2 protein from *T. repens* (A0A023HQ62, log2FC=8.9). As indicated by the name, Chlorophyll a-b binding protein (CBP) directly binds chl a/b as part of plant photosynthesis [119]. Similarly, photosystem II proteins, in particular D1/D2 and reaction center proteins, are known for their involvement in binding of pigments such as chl and cart [120–122]. As such, this substantial and significant retention in the S1 Ret can explain the visible color change and the significant drop in pigment levels (Fig. 1B) obtained in the first stage filtration. In fact, none of the proteins were reproducibly identified in S1 Perm, which is why FC is based on imputed values, further substantiating a very high level of selective retention in the first filtration stage.

Performing a similar pairwise comparison across the second filtration and subsequent diafiltration stages (S1 Perm vs. DF C), a substantially lower number of differential proteins was found (Fig. 3D). Differential analysis revealed that 84 proteins were significantly enriched in DF C while 70 proteins were depleted. This indicates a considerably lower effect on protein composition by the second filtration and DF stages compared to the first stage filtration. Surprisingly, *T. pratense* nsLTP (A0A2K3NE56), which was enriched over the first filtration stage, is significantly depleted in DF C (Log2FC=6.1) and, in fact, not identified at all. Similarly, *T. subterraneum* nsLPT (A0A2Z6P2S8) is significantly depleted (log2FC=4.4), indicating a lower allergenic risk of the final product compared to the intermediate product represented by S1 Perm. In contrast, *T. pratense* Glutathione S-transferase (GST), a known antioxidant protein [59], and several lactoylglutathione lyase isoforms (A0A2K3JKA7 and A0A2K3L521), known to participate in biosynthesis of the antioxidant glutathione [123], were enriched in the product stream, potentially improving not only the oxidative stability of the product [124,125] but also heath promoting effects [126].

Comparing the retentates and permeates from the second filtration stage (Fig. S6A) and DF (Fig. S6B), almost no enrichment in the permeate streams (2 and 4 proteins, respectively) was found. In this context, it is worth considering the underlying data normalization algorithms and missing value imputation included in LFQ analysis. This can make a direct comparison of two very different samples (e.g. the protein-rich retentates/concentrates and the dilute and protein-depleted permeates compared here) biased and may distort actual differences between samples. In contrast, a large number of proteins were enriched in the retentates (1116 and 1148 proteins, in the S2 ret and DF C respectively). Among noteworthy proteins, RuBisCO small chain from *T. repens* (P17673, log2FC=8.4 and 9.7), RuBisCO large chain from *T. subterraneum* (A0A2Z6P9S4, log2FC=9.2 and 9.3), superoxide dismutase from *T. Pratense* (A0A2K3NXG8, log2FC=3.5 and 9.5), thioredoxin M (A0A2K3P1S1, log2FC=7.4 and 8.2), and thioredoxin H (A0A2K3NRX5, log2FC=7.1 and 6.5) were highly enriched in the product streams. An enrichment of RuBisCO in the product stream is highly desirable, as RuBisCO is well acknowledged for its good nutritional and functional properties as a food protein ingredient [21–25,52]. Moreover, thioredoxins and superoxide dismutase (SOD) are known antioxidant proteins and have previously been shown to be enriched in the green juice during wet fractionation of *L. perenne*, and associated with *in vitro* antioxidant capacity of the protein extract [59].

Overall, these observations speak favorable for the product stream as a protein ingredient for food. Pigment-binding proteins are retained in the first filtration stage, improving sensory attributes of the bulk protein. Moreover, proteins with desired properties appear enriched in the product stream while minimal proteins are washed out during the second filtration stage and DF. However, as the MaxLFQ analysis uses normalized data and compares merely levels between samples, it is not evident if a differential protein is of a substantial abundance to be relevant on the bulk scale. To understand the more overall changes induced by the processing, we next performed relative in-sample quantification using relative intensity-based absolute quantification (riBAQ) to estimate the molar distribution of proteins within each stream.

### 3.5 Process-induced effects for the most abundant proteins in the streams

To identify the most abundant proteins in the respective streams and obtain a deeper insight into unit operation selectivity, we filtered the 3580 protein groups to produce a subset only containing proteins reproducibly identified (i.e. at least in two of three replicates) and with a mean relative molar abundance of at least 0.5% (by riBAQ) in any stream, excluding the unutilized side streams S2 Perm and DF Perm. Of the 3347 reproducibly identified protein groups in any stream, 63 protein groups were found above the 0.5% riBAQ threshold (Table S4). Not surprisingly, RuBisCO (rbcL and rbcS) isoforms from different species were the utmost abundant proteins by relative molar quantification using riBAQ (Fig. 4A). The high RuBisCO (rbc) levels were also reflected when summarizing all identified rbc isoforms (including those below the 0.5% threshold), where it was found to constitute around 20% of the biomass protein (Fig. 4B). High rbc levels were found in all samples upstream of DF, indicating no rbc selectivity in the first filtration stage but a high retention in the second filtration and DF stage. The high riBAQ of rbc, and particularly one rbcL isoform, in DF perm reflect identification of partially degraded protein, as these streams had very low CP and no rbc bands were detected by SDS-PAGE (Fig. 1A). As such, the content in both S2 Perm and DF Perm were not considered representative of the protein content and omitted in further analysis. An rbc content of ∼20% of the total protein in Biomass, increasing to ∼26% in Raw Juice, is somewhat lower than expected, as rbc is reported to constitute upwards of 50% of the soluble leaf protein [18,19]. In a previous study, rbc was found to constitute 35-40% of the protein in fresh green juice from *L. perenne* [59]. The slightly lower rbc content may be reflected by the somewhat lower protein yield on the analyzed batch (compared to the rest of the season) or that this particular batch had an extended (up to double of normal due to technical issues on-site) storage time (i.e. period of time from harvest to mixing with stabilizing buffer), which may facilitate partial proteolytic degradation of rbc in the green juice. Preliminary data from other batches obtained within the same season indicate that rbc content in the green juice was typically 30-50% (manuscript in preparation). Besides the high rbc levels, some general tendencies were observed among the abundant proteins across the different unit operations (Fig. 4A). While some proteins, such as Peptidyl-prolyl cis-trans isomerase (PPIase), thioredoxin-M (TRX-M), and superoxide dismutase (SOD), appear enriched on the permeate side, others, such as ATP synthase subunit β (ATPs-β), photosystem II protein D1 (PII-D1), and several chlorophyll a-b binding proteins (CBPs), become depleted. The selective depletion of particularly CBPs also correlate well with the disappearance of the ∼25 kDa band in SDS-PAGE analysis (Fig. 1A). Conversely, the opposite is observed in the S1 Ret, where the same proteins are depleted and enriched, respectively.

**Figure 4:**
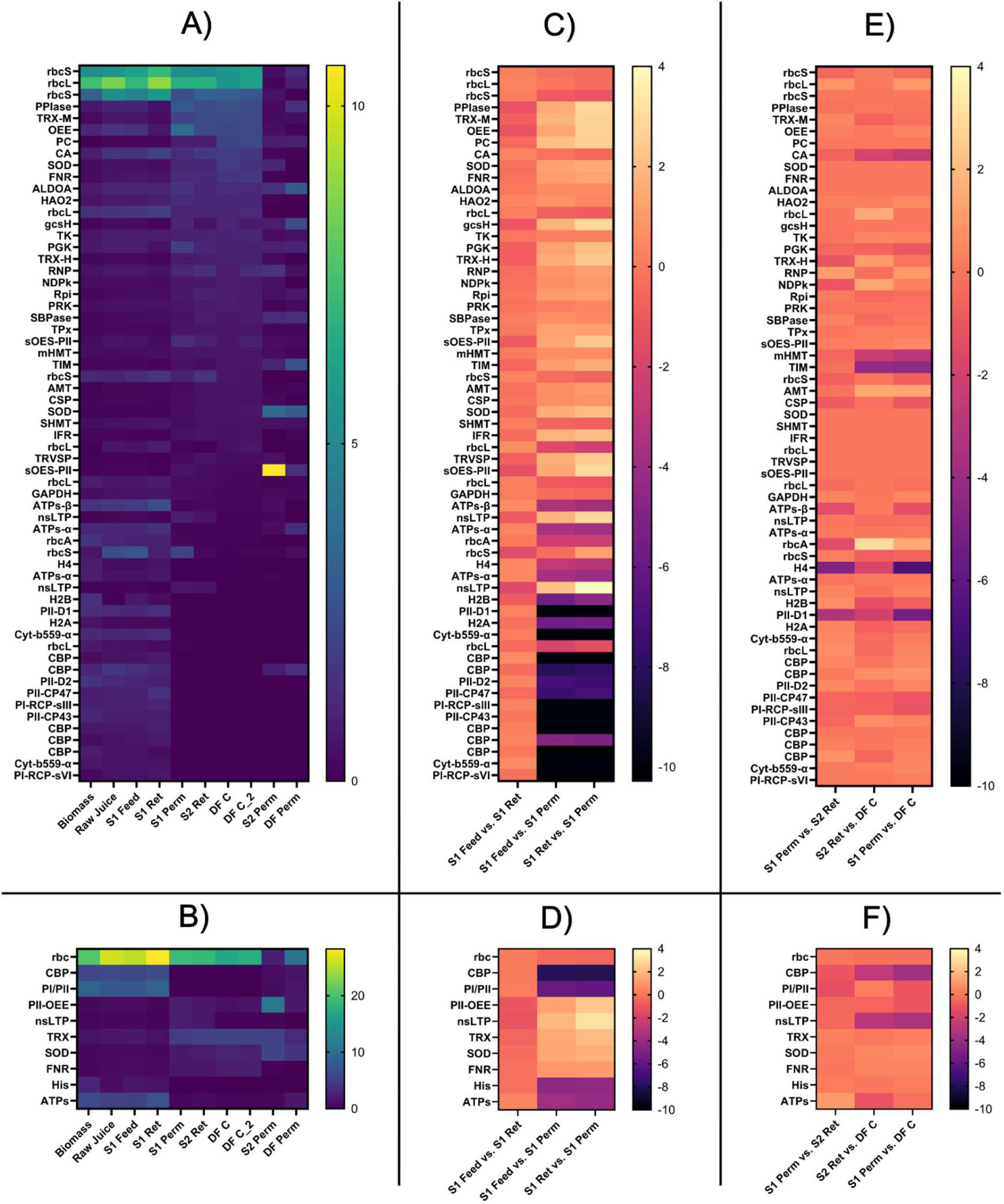
Relative molar abundance and fold-change of abundant proteins and protein families/groups. A) Heatmap representation of relative molar abundance (riBAQ) distribution for high abundance proteins (riBAQ > 0.5%) in any stream. B) Heatmap representation of total relative abundance within selected protein families/groups across all streams. C) Protein-level fold-change (by riBAQ) analysis of highly abundant proteins over the first filtration stage, represented by pair-wise comparisons of S1 Feed, S1 Ret, and S1 Perm. D) Family/group-level fold-change (by riBAQ) analysis of highly abundant protein families/groups over the first filtration stage, represented by pair-wise comparisons of S1 Feed, S1 Ret, and S1 Perm. E) Protein-level fold-change (by riBAQ) analysis of highly abundant proteins over second stage filtration and subsequent DF, represented by pair-wise comparisons of S1 Perm, S2 Ret and DF C. F) Family/group-level fold-change (by riBAQ) analysis of highly abundant protein families/groups over the second stage filtration and subsequent DF, represented by pair-wise comparisons of S1 Perm, S2 Ret and DF C. In all fold change analyses, differential abundances are represented as log2FC. If a protein (isoform or family/group) was not found or riBAQ < 0.01% in both process streams, log2FC was set to zero. For full protein names and descriptions of protein families/groups associated with the presented short names, please refer to the supplementary materials (Table S4).

To dig further into these observations, we computed the total relative abundance of all identified proteins in a grouped manner, similarly to rbc. From this analysis, the selectivity of the first stage filtration became even more evident (Fig. 4B). For certain types of proteins, such as CBPs, proteins associated with photosystem I and II (PI/II), histones (His), and ATP synthases (ATPs), a high level of selective retention was observed. Next, we performed fold-change (FC) analysis across the first filtration stage, where S1 Feed, S1 Ret, and S1 Perm were compared in pair-wise FC analysis. In this analysis, the riBAQ of both individual proteins (Fig. 4C) and grouped proteins (Fig. 4D) were used. The FC analysis further substantiates the depletion of the aforementioned proteins and groups across the first stage filtration, and in fact, CBPs were depleted from 5.9% in S1 Feed to <0.03% in S1 Perm while PI/II were depleted from 7.6% to 0.1%. The specific depletion of these proteins is considered crucial for removing the undesired color due to their ability to bind pigments such as chlorophylls and carotenoids [127–129]. In contrast, proteins with known antioxidant properties such as thioredoxins (TRX), SODs and ferredoxins (FNR) become enriched following first stage filtration from respectively 1.7%, 0.7% and 0.8% in S1 Feed (Fig. 4B) to 4.6%, 1.8%, and 1.6% in S1 perm, representing a 2.8, 2.6, and 1.9-fold enrichment (Fig. 4D). Similarly, nsLTPs were enriched 3.8-fold while the oxygen evolving proteins of photosystem II (PII-OES), in contrast to pigment binding PI/II proteins, were also enriched 2.5-fold.

A much less pronounced effect on the quantitative protein composition was observed across the second filtration and DF stages (Fig. 4E). Most notably, plastocyanin (PC) and some rbcL isoform were enriched while other rbcL, rbcS, and rbcA isoforms were depleted. Several nsLPT isoforms, which were enriched in the permeate from the first stage filtration (S1 Perm), were depleted substantially (8- to 20-fold) while any residual CBPs were also depleted. Using the protein groupings (Fig. 4F), CBPs, which were substantially depleted in the first stage filtration, were a further 14-fold depleted to less than 0.002% in DF C from an initial level of 5.9% in S1 Feed. This represents a reduction of >99.9% of CBPs across the entire process. Similarly, PI/II proteins were further depleted over the second filtration and DF stages, resulting in a final abundance of 0.06% in DF C from an initial abundance of 7.6% in S1 Feed, thereby representing a removal of > 99% across the whole process. The total abundance of nsLTPs was depleted to a final abundance of 0.21% from 2.15% in S1 Perm, thereby representing a removal of >90% of the potential allergen, which was enriched in the first filtration stage. Based on this observation, it is not unlikely that nsLTPs can be further depleted by extending the DF stage to allow more washing of the protein. Reticuline oxidase-like protein (A0A2K3N926), which was found enriched in S1 Perm by LFQ-based analysis, was generally found of low abundance across all streams (< 0.1%) and found in very low abundance (< 0.004%) in the final product (DF C). As such, this allergen is not considered of particular relevance at the bulk level.

### 3.6 Process-induced changes cause notable changes to mean protein properties

Next the mean physicochemical properties of the process stream proteins were investigated using *in silico* analysis. An unweighted analysis was initially performed based only on reproducibly identified proteins in the different process streams. For this, a range of physicochemical, allergenic, subcellular, and *in silico* structural properties were computed for all identified proteins. The mean property within each stream was determined by computing the mean for all proteins identified in that process stream. This facilitated investigation of any enrichment or protein-level clustering based on membrane selectivity in the different unit operations. Using computed physicochemical and structural properties as descriptors, no substantial differences between the different streams were observed (Fig. S7A-H) despite the fact that isoelectric point (pI) and structural features were found to be significant (p<0.05) in explaining data variability by ANOVA (Table S5). No significant clustering was observed based on either the protein occurrence in specific streams (Fig. S8A) or by their subcellular localization (Fig. S8B) using TruncatedSVD-based dimensionality reduction.

As unweighted analysis did not provide any noteworthy insights, the protein-level abundances by riBAQ were used as a weights for calculating the mean properties within each process streams. This approach proved to be much more effective for describing differential mean properties across the different process streams. For instance, distinct differences in mean properties were found between the different process streams (Fig. S9A-H). In fact, all properties were found to be highly significant (p<1E-6) for describing data variability by one-way ANOVA (Table S6). These differences were also reflected by distinct clustering using TruncatedSVD-based dimensionality reduction from computed physicochemical and structural properties (Fig. S10). Based on physicochemical and structural properties, samples upstream of the first filtration stage (including S1 Ret) and samples downstream clustered nicely while the two unutilized side streams (S2 Perm and DF Perm) were substantially more scattered. Due to the low CP and number of protein IDs, these streams were omitted from subsequent analysis. As a result, two highly distinct clusters were formed (Fig. 5A), an “upstream cluster” and a “downstream cluster” based on their origin relative to the first filtration stage. The same clustering of streams was observed using predicted subcellular localization (Fig. 5B), highlighting the distinct differences in quantitative protein composition.

**Figure 5:**
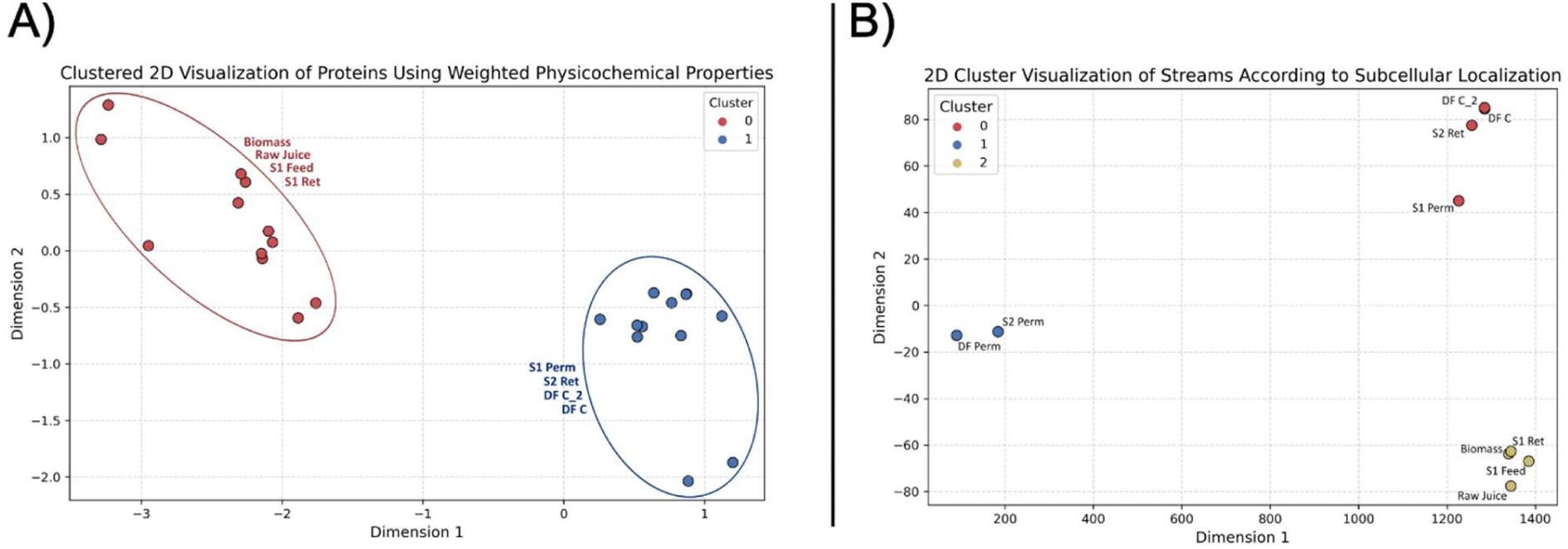
Two-dimensional representation of protein across process streams obtained by TruncatedSVD, followed by clustering. A) riBAQ-weighted computed physicochemical and structural properties (replicate level) with unutilized waste streams (S2 Perm and DF Perm) omitted. B) riBAQ-weighted subcellular localization (sample level) across all process streams.

To identify which properties are important for the observed clustering, the distribution of each property across clusters was visualized using boxplots of the eight descriptive properties individually (Fig 6A-H). Across all eight properties, distinct and significant (p<0.05) differences were found between the upstream cluster (cluster 0) and the downstream cluster (cluster 1). Upstream samples were found to have larger median molecular weight, aromaticity, instability index, GRAVY, and pI. Upstream samples were in general also found to have a larger proportion of well-defined secondary structure with a higher content of both α-helix and β-sheet while the proportion of turn were decreased.

**Figure 6:**
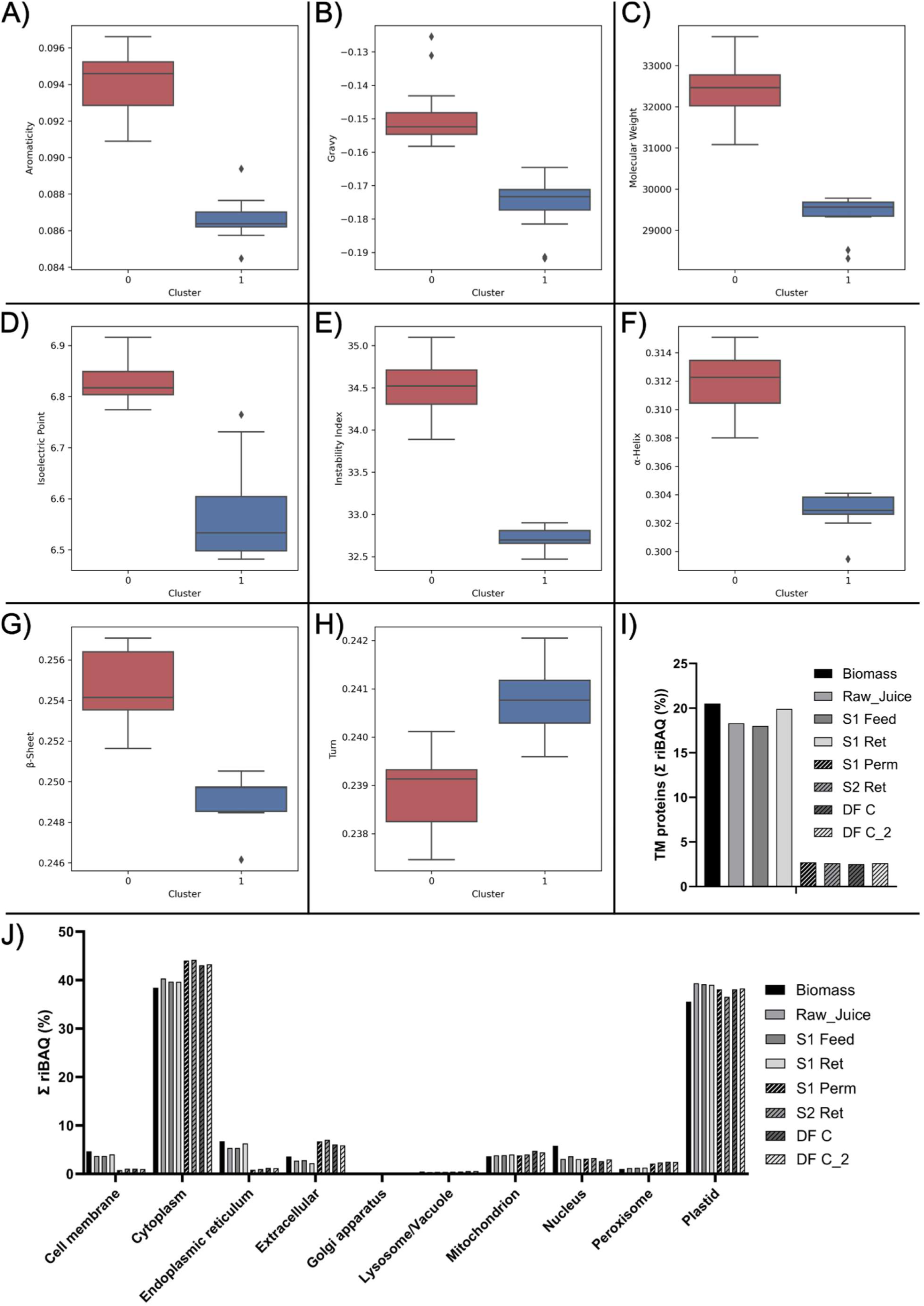
Comparison of riBAQ-weighted cluster properties and subcellular distribution. A-H) Boxplots of riBAQ-weighted computed properties of the two process stream clusters (using TruncatedSVD dimensionality reduction) shown for aromaticity (A), GRAVY (B), molecular weight (C), isoelectric point (D), instability index (E), α-helical fraction (F), β-sheet fraction (G), and turn fraction (H). For all properties, differences between clusters were significant with p < 0.0002. I) riBAQ-weighted abundance of proteins with transmembrane regions, predicted with TOPCONS2, within each process stream. J) riBAQ-weighted subcellular distribution, predicted using DeepLoc, of proteins within each process stream. For transmembrane regions and protein subcellular localization, the mean riBAQ (n=3) for the individual streams were plotted.

The higher abundance of membrane-associated proteins in upstream samples, as indicated by pair-wise comparison of S1 Ret and S1 Perm using LFQ data (Fig. 3E-F), correlates well with the differences in physicochemical properties between the clusters. Plant membrane proteins are similarly reported to generally have a higher MW and more basic pI compared to soluble proteins [130], which correlates with a higher median MW and pI found in the upstream cluster (Fig. 6C-D). Membrane proteins contain transmembrane domains and often are responsible for pore formation [131], why a closer to zero GRAVY in the upstream cluster (Fig. 6B) reflects their more amphiphilic nature [132]. While reduced amphiphilicity may reduce the ability of the final protein product to e.g. form stable oil-in-water emulsions [133], the larger negative GRAVY also indicates a better solubility [134]. These agree with recent work on the functionality of a protein product produced using the presented membrane-based biorefinery concept, where a high solubility was observed while less impressive emulsifying properties were seen [52]. As membrane proteins require a highly ordered structure, particularly in membrane-spanning regions [131,132], this also explains the higher proportion of defined secondary structure elements (Fig. 6F-H). Furthermore, membrane proteins are reported to have a lower stability [131], correlating well with a higher instability index in the upstream cluster (Fig. 6E), particularly prominent in the TM-enriched S1 Ret (Fig. 3F, Fig. 6I, Fig. S9E).

Considering the differences in subcellular localization of proteins, subtle but still distinct differences were observed when performing unweighted analysis (Table S7 and Table S8). A larger proportion of membrane proteins were found in upstream samples, particularly in S1 Ret, in agreement with the pairwise comparison of S1 Ret and S1 Perm (Fig. 3E). Similarly, upstream samples generally contained higher proportions of proteins localized in the ER, Golgi, and mitochondria, while downstream proteins contained a higher proportion of cytoplasmic and extracellular proteins. These differences were more pronounced when factoring relative protein abundances (riBAQ) in a weighted analysis (Fig. 6J). Here, a selective retention of cell membrane and ER (the site for transmembrane protein synthesis [135]) proteins were observed across the first filtration stage. Cell membrane proteins constituted 3.7% of the total protein in S1 Feed but were depleted to 0.8% in S1 Perm. In contrast, the proportion of cytoplasmic and extracellular protein increased. Only subtle differences were observed when comparing S1 Perm, S2 Ret and the final product DF C. The selective retention of membrane-associated proteins is further substantiated when considering the distribution of all proteins predicted to comprise transmembrane regions (Fig. 6I). Here, the abundance of TM proteins in S1 Feed (18%) was substantially depleted over the first membrane into S1 Perm (2.7%) and slightly enriched in S1 Ret (20%). These results corroborated the findings of the LFQ-based pair-wise comparison between S1 Ret and S1 Perm (Fig. 3E-F). In the final product stream (DF C), the low level of membrane-associated proteins was at a similar level as in S1 Perm (2.5%), representing a 7.2-fold depletion of membrane-associated proteins compared to S1 Feed. Ultimately, our findings show a high degree of selectivity towards particularly membrane-associated and known pigment-binding proteins in the first filtration stage of the biorefinery process, being responsible for significant changes in optical properties and sensory attributes of the different streams. Meanwhile, the second filtration and DF stages, using a 10 kDa cutoff membrane, merely retains and concentrates proteins in a non-discriminatory manner while washing out small molecule contaminants, phytochemicals, and residual pigments.

## 4. Conclusions

Recently, a nondestructive two-stage membrane-based biorefinery concept was developed to produce native leaf protein devoid of grassy attributes. However, the underlying reasons why this process works remain unexplored. In this study, process streams from each unit operation within the two-stage membrane-process were characterized to understand selectivity and driving forces from a molecular perspective. The process efficiently removes >99.9% of investigated pigments (chlorophylls and carotenoids), with the majority being retained in the first stage filtration. Using mass spectrometry-based proteomics with two complementary quantification strategies and subsequent bioinformatic analysis, this was found to be correlated with a selective retention of membrane-associated and pigment-binding proteins, particularly in the first filtration stage. Over the full process, >99.9% of chlorophyll-binding proteins and other pigment-binding proteins from photosystems I and II were removed from the final protein concentrate, obtained following a second filtration and subsequent diafiltration (DF) stage. In contrast, proteins with reported antioxidant properties were enriched in the final protein product intended for use as a food ingredient, which may improve not only stability of the product but also potential health benefits from ingestion. Moreover, proteins with known allergenic properties were found to be depleted by the biorefinery concept. Importantly, a high abundance of RuBisCO was found to be retained throughout the process, explaining the promising properties of the protein concentrate, previously described.

This study clearly demonstrates that the larger pore membrane used in first-stage filtration displays a much higher degree of differential protein retention compared to 10 kDa membrane used in the second filtration and DF stages. This also reflects why both stages were essential for a successful process, as the first stage initially retains undesired large components. Subsequently, the second stage and DF facilitates permeation of residual small molecule contaminants (e.g. free pigments and other phytochemicals) as well as buffer components and salts from the stabilizing solution employed in the process. This stage facilitates concentrating the protein to a higher purity, while retaining the quantitative composition, and ultimately producing a final product stream for subsequent drying. Overall, this study provides a molecular perspective and protein-level insights on membrane selectivity in green biorefineries and highlights what proteins should be selectively removed from green juice to improve quality and functionality of a protein product.

## Data availability

The majority of data is available in the supplementary information. The mass spectrometry proteomics data have been deposited into the ProteomeXchange Consortium via the PRIDE [83] partner repository with the dataset identifier PXD069787 and 10.6019/PXD069787. All other data can be made available upon request.

## Author contributions

SGE: Conceptualization, Methodology, Validation, Formal analysis, Investigation, Visualization, Funding acquisition, Project administration, Supervision, Writing – original draft preparation, Writing – review and editing. NAK: Methodology, Formal analysis, Investigation, Visualization, Writing – review and editing. NHJ: Formal analysis, Investigation, Writing – review and editing. AKJ: Resources, Formal analysis, Investigation, Writing – review and editing. TM: Resources, Investigation, Writing – review and editing. MKJ: Resources, Investigation, Writing – review and editing. PSL: Conceptualization, Funding acquisition, Project administration, Writing – review and editing. ML: Conceptualization, Funding acquisition, Project administration, Supervision, Writing – review and editing.

## Declaration of competing interest

Peter Stephensen Lübeck reports a relationship with BiomassProtein ApS that includes board membership. Simon Gregersen Echers, Peter Stephensen Lübeck, Tuve Mattsson, Anders K Jørgensen, Mads K Jørgensen, and Mette Lübeck have the patent #WO2025/133209: Method for Producing a food-grade protein product and/or feed protein product from plant material. Remaining authors report no competing interests.

## Funding

This work was supported by the Danish Green Development and Demonstration Program (GUDP) under the “Grass4Food” project [grant number 34009-20-1657] and the Independent Research Foundation Denmark (DFF) under the “PhyPro” project [grant number DFF 1127-00245B].

## Acknowledgements

The authors would like to acknowledge the laboratory assistance provided by laboratory technician Sofie Albrekt Hansen.

## Notes

### Competing Interest Statement

Peter Stephensen Lubeck reports a relationship with BiomassProtein ApS that includes board membership. Simon Gregersen Echers, Peter Stephensen Lubeck, Tuve Mattsson, Anders K Jorgensen, Mads K Jorgensen, and Mette Lubeck have the patent #WO2025/133209: Method for Producing a food-grade protein product and/or feed protein product from plant material. Remaining authors report no competing interests.

