## Supplementary Information for "Selective removal of green pigments and associated proteins from clover-grass protein concentrates: Molecular insight into a non-destructive, two-stage membrane-based biorefinery concept for high-quality food protein production"

**For**

**Note to the reader:**

In the raw data supplied (e.g. LC-MS/MS) a different nomenclature is applied to reflect the internal, daily naming of the process steps. As such, samples from first stage filtration (S1) are referred to as microfiltration (MF) while the use of a 60nm membrane does not qualify as microfiltration per definition but is in principle ultrafiltration. Similarly, the second stage filtration (S2) is referred to as ultrafiltration (UF) despite both stages in principle represent ultrafiltration. The final product stream (DF C) may be referred to as "DC" while its replicate (DF C_2) may be referred to as "DF Ret". This discrepancy in sample naming is reflected in some plots within the supplementary material.

### Content of supplementary material

#### Supplementary tables

**Table S1:** MaxQuant output data and additional downstream processing hereof.

**Table S2:** Uniprot AC#s for protein families/types/groups

**Table S3:** Differential proteins from LFQ-based pair-wise comparison of MF Ret and MF Perm.

**Table S4:** Abundant proteins (riBAQ > 0.5%) across all streams.

**Table S5:** One-Way ANOVA for unweighted analysis using computed physicochemical properties.

**Table S6:** One-Way ANOVA for riBAQ-weighted analysis using computed physicochemical properties.

**Table S7:** Distribution of protein subcellular localization as predicted by DeepLoc (by count)

**Table S8:** Distribution of protein subcellular localization as predicted by DeepLoc (by %)

#### Supplementary figures

**Figure S1:** One-way ANOVA of dry matter and crude protein across streams.

**Figure S2:** Quantification of individual pigments relative to dry matter and crude protein.

**Figure S3:** Optimization of protein extraction.

**Figure S4:** Distribution of protein ID's and sample similarity analysis (replicate level).

**Figure S5:** Pair-wise differential analysis of MF Ret vs. MF Perm and overlap with differential proteins from pair-wise comparison of MF Feed vs. MF Perm.

**Figure S6:** Pair-wise differential analysis of retentates and permeates from UF and DF.

**Figure S7:** Boxplots showing the distribution of computed physicochemical properties by sample based on unweighted analysis.

**Figure S8:** Two-dimensional representation of proteins obtained via TruncatedSVD from unweighted computed physicochemical properties by sample and predicted subcellular localization.

**Figure S9:** Boxplots showing the distribution of computed physicochemical properties by sample based on riBAQ-weighted analysis.

**Figure S10:** Two-dimensional representation of protein properties from each stream (replicate-level) obtained via TruncatedSVD on riBAQ-weighted computed physicochemical properties.

**Table S1:** MaxQuant output data and additional downstream processing hereof. The "proteinGroups" folder has been modified to include a range of additional data used in the manuscript covering calculation of relative iBAQ (riBAQ), replicate means, abundance and reproducibility filters. For a full data file with no modification, see the referenced project in the PRIDE data repository. Table can be found in appended .xlsx file (Supplementary Table) as "Table S1".

**Table S2:** Uniprot AC#s for protein families/types/groups. For each defined family/type/group, the AC# of the lead protein (i.e. first protein within each MaxQuant proteinGroup) is listed. Table can be found in appended .xlsx file (Supplementary Table) as "Table S2".

**Table S3:** Differential proteins from LFQ-based pair-wise comparison of MF Ret and MF Perm. All differentially abundant proteins are included and the list also includes fold change (as log2FC) and associated p-value from Mass Dynamics. For all proteins, the predicted subcellular localization (by DeepLoc) and the inclusion of probable transmembrane (TM) regions (by TOPCONS2) are included in the table along with lead protein AC#s and protein names. Table can be found in appended .xlsx file (Supplementary Table) as "Table S3".

**Table S4:** Overview of the most abundant (riBAQ > 0.5%) proteins in any of the analyzed samples. The table includes the Uniprot AC#, protein name, a short name (as used in Figure 4), various MaxQuant Data (number of proteins in group, peptide IDs, unique peptides, sequence coverage, molecular weight and Andromeda score), mean riBAQ abundance across all sample types, and log2FC (by riBAQ abundance) from various comparisons (see headers). For proteins not detected in a particular stream, log2FC was set to +/- 10 based on maximal values observed in the analysis. If a protein was not found or riBAQ < 0.01% in both streams, log2FC was set to zero. The table contains both individual proteins (MaxQuant proteinGroups) and the defined "families/types/groups" from this work (see Table s2). Table can be found in appended .xlsx file (Supplementary Table) as "Table S4".

**Table S5:** Results from One-Way ANOVA of computed physicochemical properties across streams based on unweighted analysis.

| Property | ANOVA F | ANOVA p-value |
|---|---|---|
| MW | 0.83 | 0.56 |
| Aromaticity | 2.00 | 0.05 |
| Instability Index | 0.39 | 0.91 |
| GRAVY | 0.89 | 0.51 |
| pI | 19.45 | 0.00 |
| Alpha Helix fraction | 2.62 | 0.01 |
| Turn fraction | 6.11 | 0.00 |
| Beta Sheet fraction | 3.60 | 0.00 |

**Table S6:** Results from One-Way ANOVA of computed physicochemical properties across streams based on riBAQ-weighted analysis.

| Property | ANOVA F | ANOVA p-value |
|---|---|---|
| MW | 85.19 | 1.82E-11 |
| Aromaticity | 39.73 | 6.02E-09 |
| Instability Index | 105.87 | 3.37E-12 |
| GRAVY | 36.41 | 1.15E-08 |
| pI | 22.01 | 4.39E-07 |
| Alpha Helix fraction | 98.98 | 5.69E-12 |
| Turn fraction | 12.86 | 1.66E-05 |
| Beta Sheet fraction | 27.46 | 9.04E-08 |

**Table S7:** Distribution of DeepLoc-predicted subcellular localizations across streams (by count).

| Process Stream | Cell membrane | Cytoplasm | Endoplasmic reticulum | Extracellular | Golgi apparatus | Lysosome/Vacuole | Mitochondrion | Nucleus | Peroxisome | Plastid |
|---|---|---|---|---|---|---|---|---|---|---|
| Biomass | 77 | 1117 | 124 | 193 | 28 | 60 | 200 | 138 | 58 | 653 |
| Raw_Juice | 70 | 1111 | 117 | 189 | 27 | 63 | 187 | 136 | 65 | 681 |
| S1 Feed | 76 | 1148 | 117 | 203 | 26 | 66 | 190 | 148 | 66 | 691 |
| S1 Ret | 82 | 1122 | 118 | 189 | 27 | 68 | 188 | 137 | 64 | 659 |
| S1 Perm | 39 | 1051 | 66 | 230 | 18 | 54 | 124 | 118 | 61 | 555 |
| S2 Ret | 49 | 1092 | 68 | 246 | 18 | 58 | 125 | 126 | 65 | 534 |
| DF C | 51 | 1118 | 81 | 261 | 18 | 56 | 130 | 127 | 66 | 540 |
| DF C_2 | 44 | 1118 | 78 | 259 | 19 | 62 | 131 | 128 | 68 | 540 |
| S2 Perm | 9 | 148 | 3 | 29 | 0 | 3 | 15 | 10 | 13 | 107 |
| DF Perm | 4 | 71 | 0 | 8 | 0 | 1 | 9 | 6 | 6 | 58 |

Table S8. Distribution of DeepLoc-predicted subcellular localizations across streams (by % of all reproducibly identified proteins within the stream).

| Process Stream | Cell membrane | Cytoplasm | Endoplasmic reticulum | Extracellular | Golgi apparatus | Lysosome/Vacuole | Mitochondrion | Nucleus | Peroxisome | Plastid |
|---|---|---|---|---|---|---|---|---|---|---|
| Biomass | 2.9% | 42.2% | 4.7% | 7.3% | 1.1% | 2.3% | 7.6% | 5.2% | 2.2% | 24.7% |
| Raw_Juice | 2.6% | 42.0% | 4.4% | 7.1% | 1.0% | 2.4% | 7.1% | 5.1% | 2.5% | 25.7% |
| S1 Feed | 2.8% | 42.0% | 4.3% | 7.4% | 1.0% | 2.4% | 7.0% | 5.4% | 2.4% | 25.3% |
| S1 Ret | 3.1% | 42.3% | 4.4% | 7.1% | 1.0% | 2.6% | 7.1% | 5.2% | 2.4% | 24.8% |
| S1 Perm | 1.7% | 45.4% | 2.8% | 9.9% | 0.8% | 2.3% | 5.4% | 5.1% | 2.6% | 24.0% |
| S2 Ret | 2.1% | 45.9% | 2.9% | 10.3% | 0.8% | 2.4% | 5.2% | 5.3% | 2.7% | 22.4% |
| DF C | 2.1% | 45.7% | 3.3% | 10.7% | 0.7% | 2.3% | 5.3% | 5.2% | 2.7% | 22.1% |
| DF C_2 | 1.8% | 45.7% | 3.2% | 10.6% | 0.8% | 2.5% | 5.4% | 5.2% | 2.8% | 22.1% |
| S2 Perm | 2.7% | 43.9% | 0.9% | 8.6% | 0.0% | 0.9% | 4.5% | 3.0% | 3.9% | 31.8% |
| DF Perm | 2.5% | 43.6% | 0.0% | 4.9% | 0.0% | 0.6% | 5.5% | 3.7% | 3.7% | 35.6% |

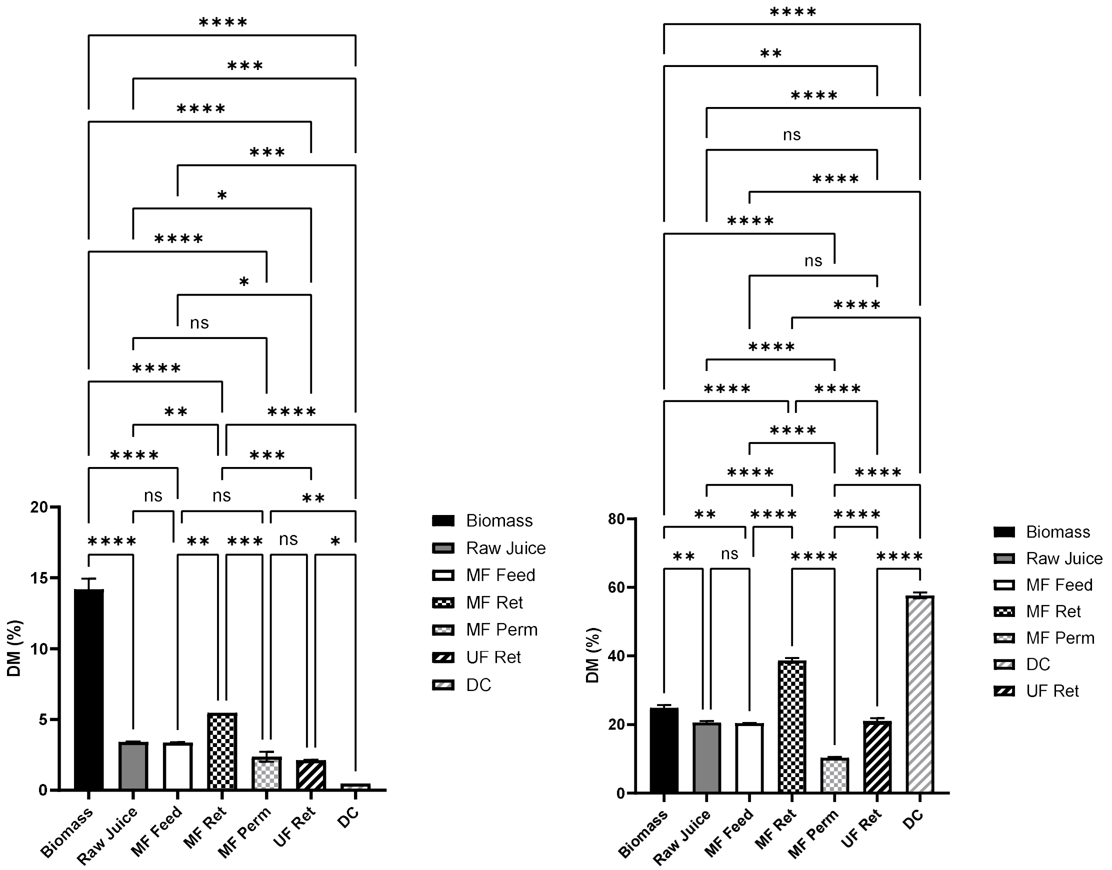


**Figure S1:** One-way ANOVA of dry matter (left) and crude protein (right) across streams. Analysis was performed using Tukey with a 95% confidence interval. Values are indicated as means with the standard deviation (2). Statistical analysis is performed as one-way ANOVA with significance level (from adjusted p-values) indicated by "ns" ($p > 0.05$), "*" ($p \leq 0.05$), "**" ($p \leq 0.01$), "***" ($p \leq 0.001$), and "****" ($p \leq 0.0001$).

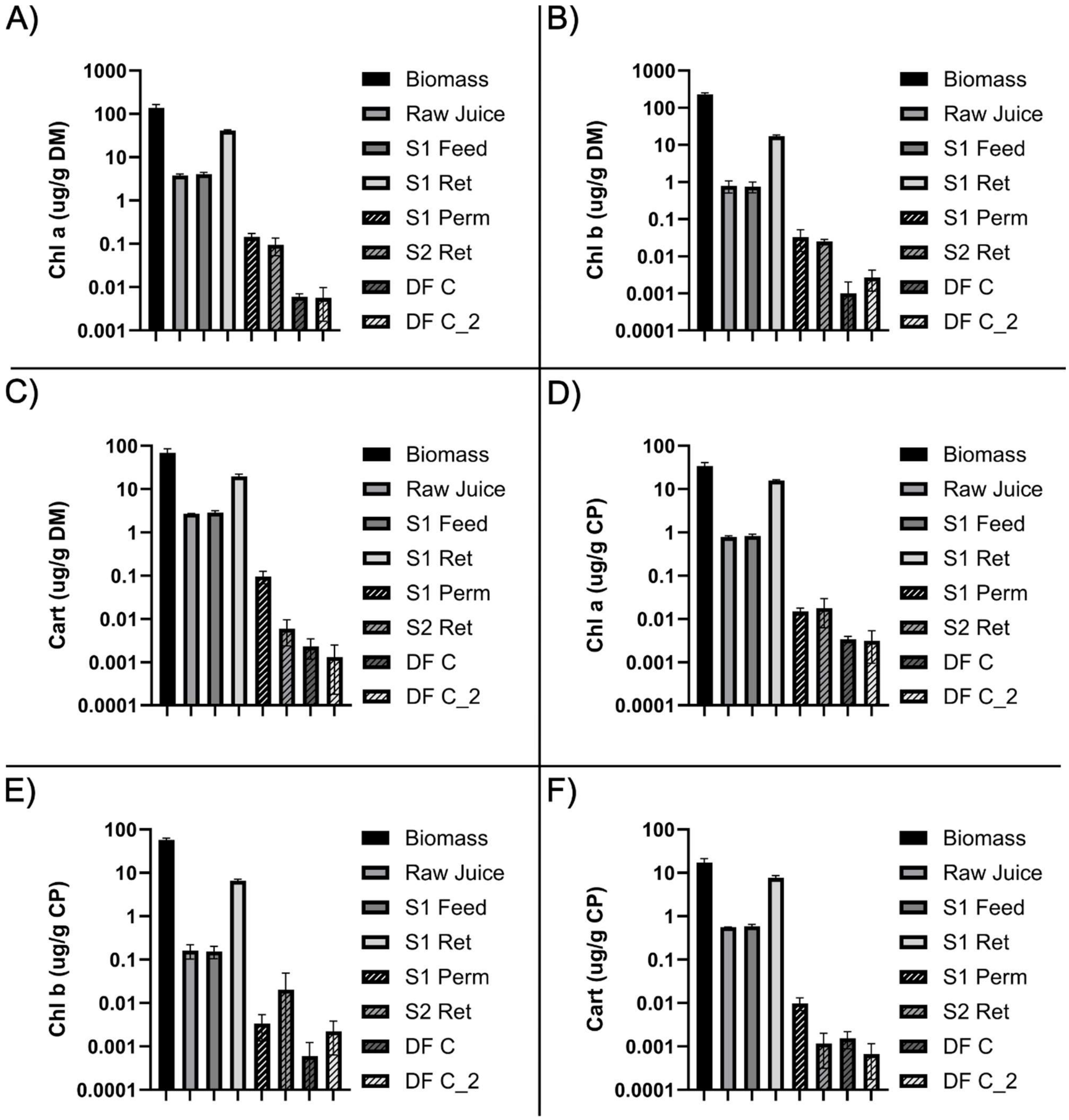


**Figure S2**: Quantification of individual pigments by UV/Vis. Pigment content is presented relative to DM for Chlorophyll-a (A), Chlorophyll-b (B) and Total Carotenoids (C). Pigment content relative to CP for Chlorophyll-a (D), Chlorophyll-b (E), and total Carotenoids (F). Note that results are presented using a Log10 scale.

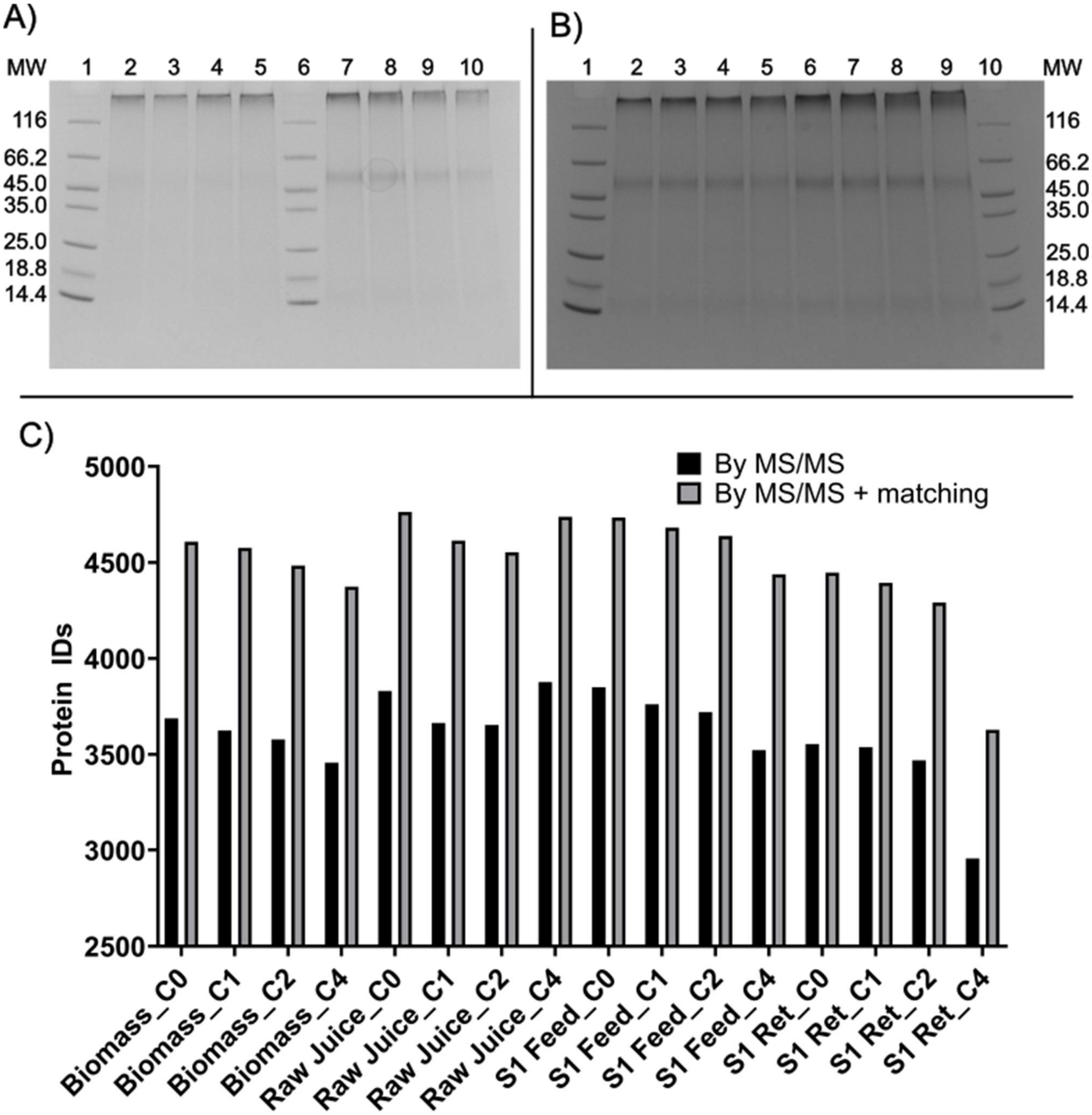


**Figure S3:** Optimization of protein extraction. A) Reducing SDS-PAGE of AFA (0,1,2,4 cycles) from unprocessed biomass (lane 2-5) and raw green juice (lane 7-10). MW marker is included in lane 1 and 6. B) Reducing SDS-PAGE of AFA (0,1,2,4 cycles) from S1 Feed (lane 2-5) and S1 Ret (lane 6-9). MW marker is included in lane 1 and 10. C) Number of peptide IDs for the different streams using increasing number of AFA cycles (CX) where X represents the number of cycles. Both the number of peptide IDs from MS/MS alone and with "matching between runs" is shown.

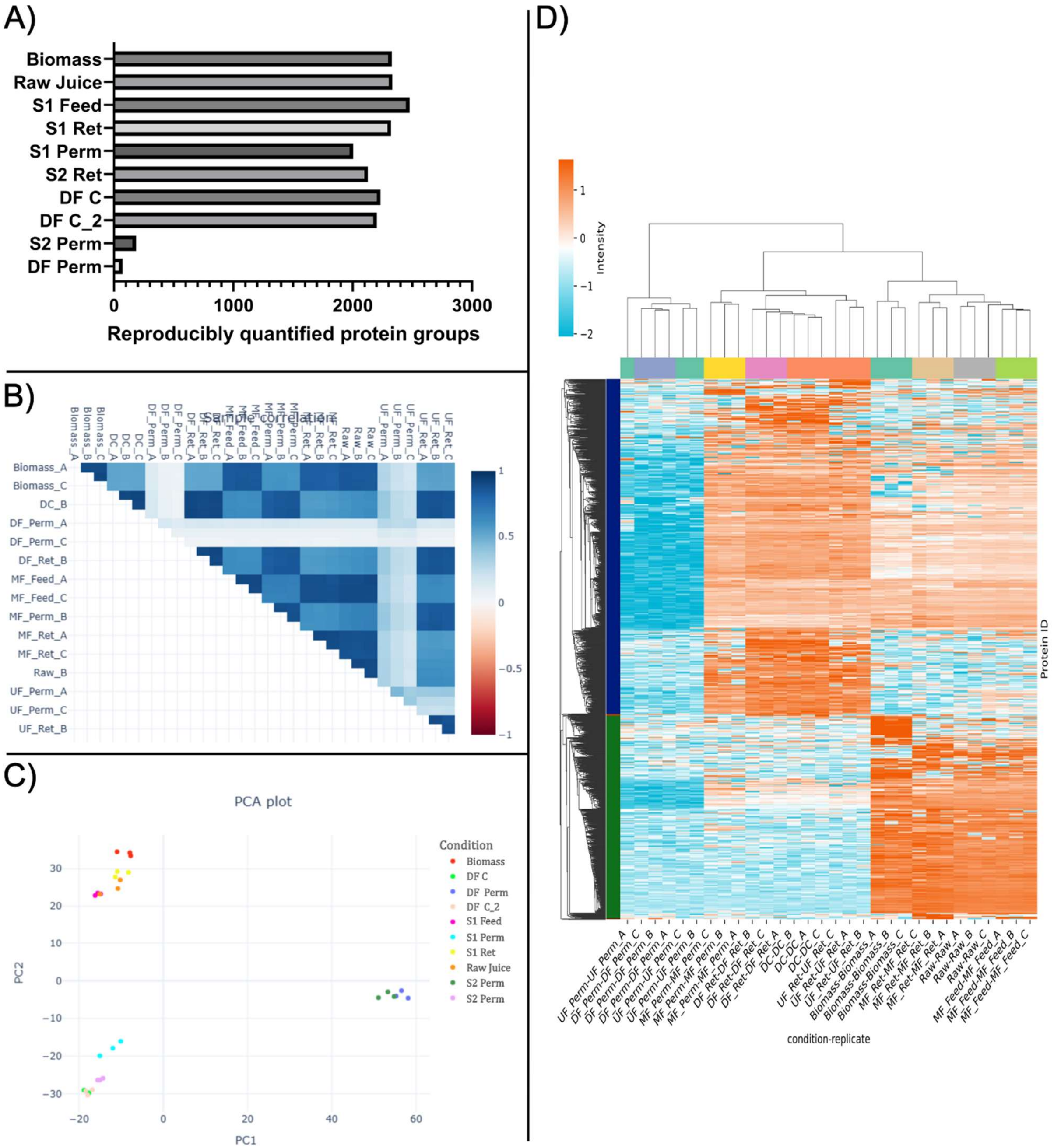


**Figure S4:** Distribution of protein ID's and sample similarity analysis (replicate level). A) Reproducibly quantified (i.e. quantified in at least two of three replicates of each stream) proteins across all streams. B) Pair-wise sample similarity (replicate-level) by Pearson Correlation Coefficient (PCC) analysis. C) Two-dimensional representation of sample similarity by PCA for MaxLFQ-based quantification in Mass Dynamics. D) Heatmap representation of differentially abundant proteins by ANOVA analysis in MassDynamics. Data is depicted as z-score standardized MaxLFQ intensities by row (protein group) and clustered using a Euclidian distance of 7.

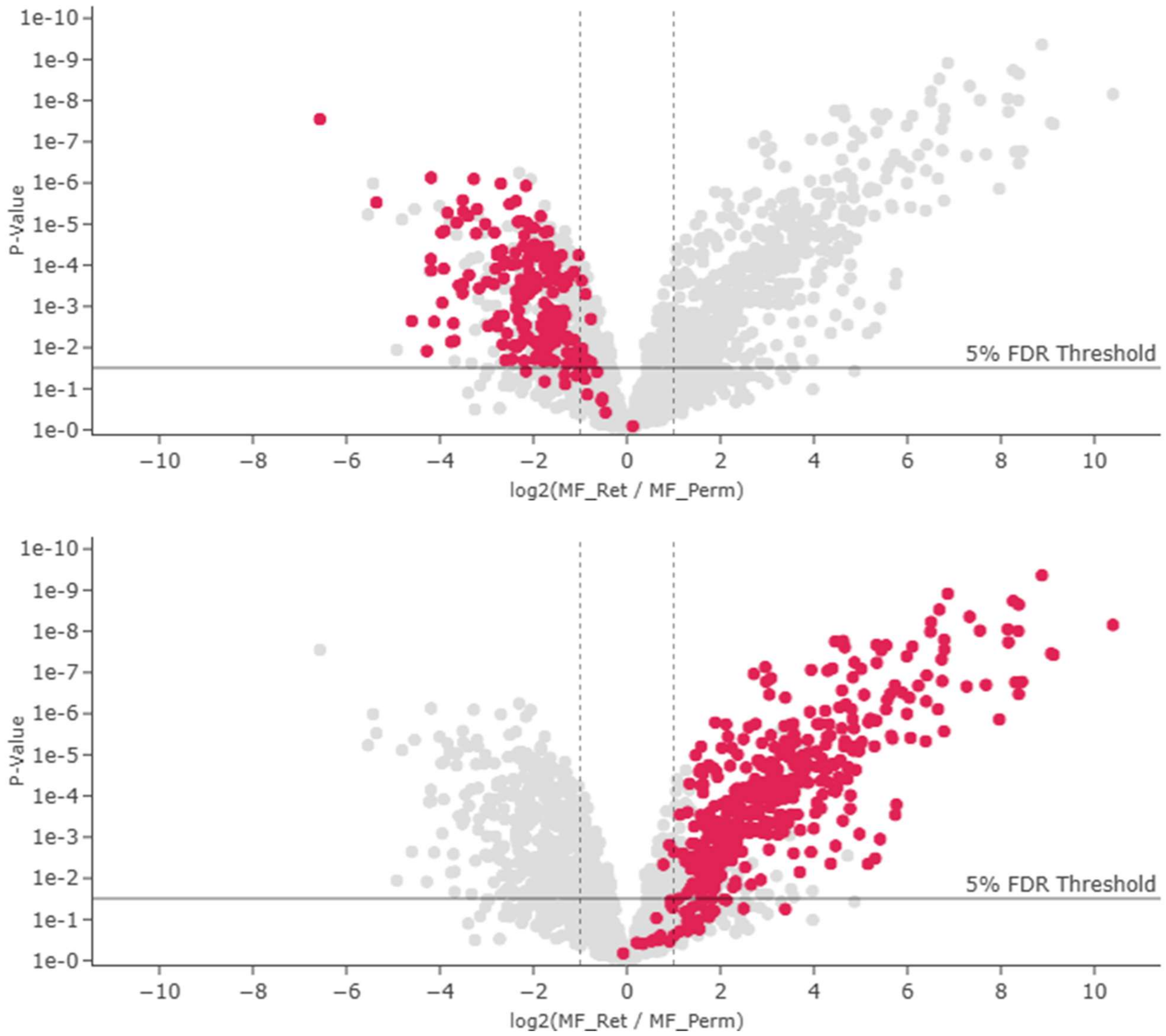


**Figure S5:** Pair-wise differential analysis of MF Ret vs. MF Perm and overlap with differential proteins from pair-wise comparison of MF Feed vs. MF Perm. Volcano plots from pair-wise analysis of MF Ret vs. MF Perm with overlay (highlighted in red) of proteins enriched in MF Feed (top) and in MF Perm (bottom) from the pair-wise comparison of MF Feed vs. MF Perm (Fig. 3C). Analysis and plots performed in Mass Dynamics using MaxLFQ-based quantification.

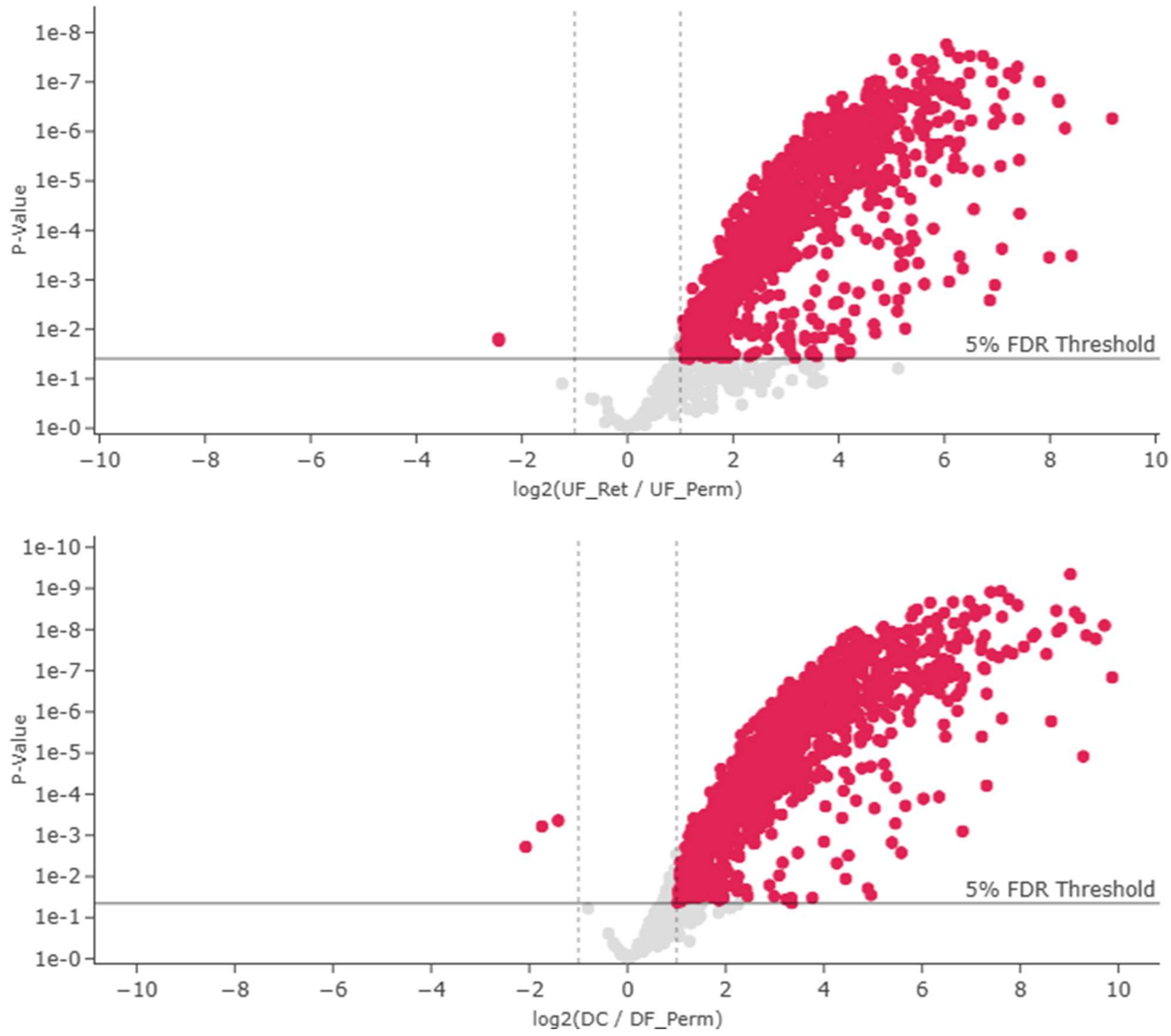


**Figure S6:** Pair-wise differential analysis of retentates and permeates from UF and DF. Volcano plot from pair-wise comparison of UF Ret vs. UF Perm (top) and DC vs. DF Perm (bottom). Differentially abundant proteins are highlighted in red. Analysis and plots performed in Mass Dynamics using MaxLFQ-based quantification.

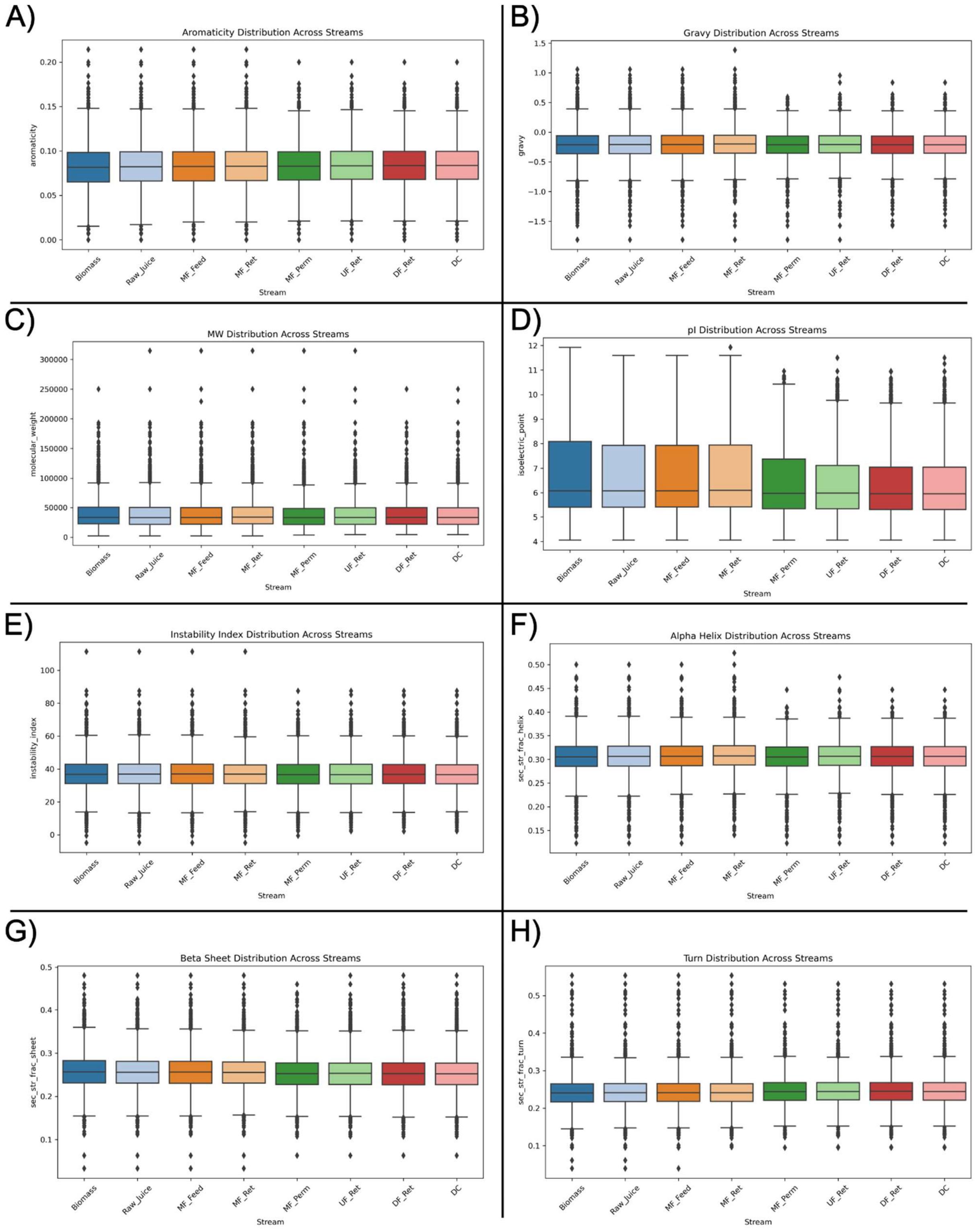


**Figure S7:** Boxplots showing the distribution of computed physicochemical properties by stream (S2 Perm and DF Perm excluded) based on unweighted analysis. Properties computed cover: Aromaticity (A), GRAVY (B), molecular weight (C), isoelectric point (D), instability index (E), α-helical content (F), β-sheet content (G), and turn content (H).

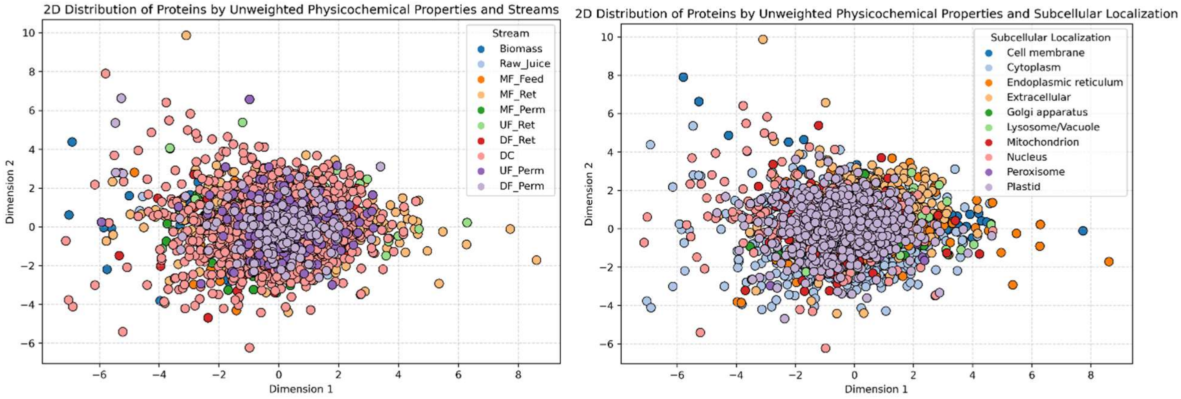


**Figure S8:** Two-dimensional representation of proteins obtained via TruncatedSVD from unweighted computed physicochemical properties by sample (left) and predicted subcellular localization (right). Each dot represents a protein in the respective stream or predicted subcellular localization.

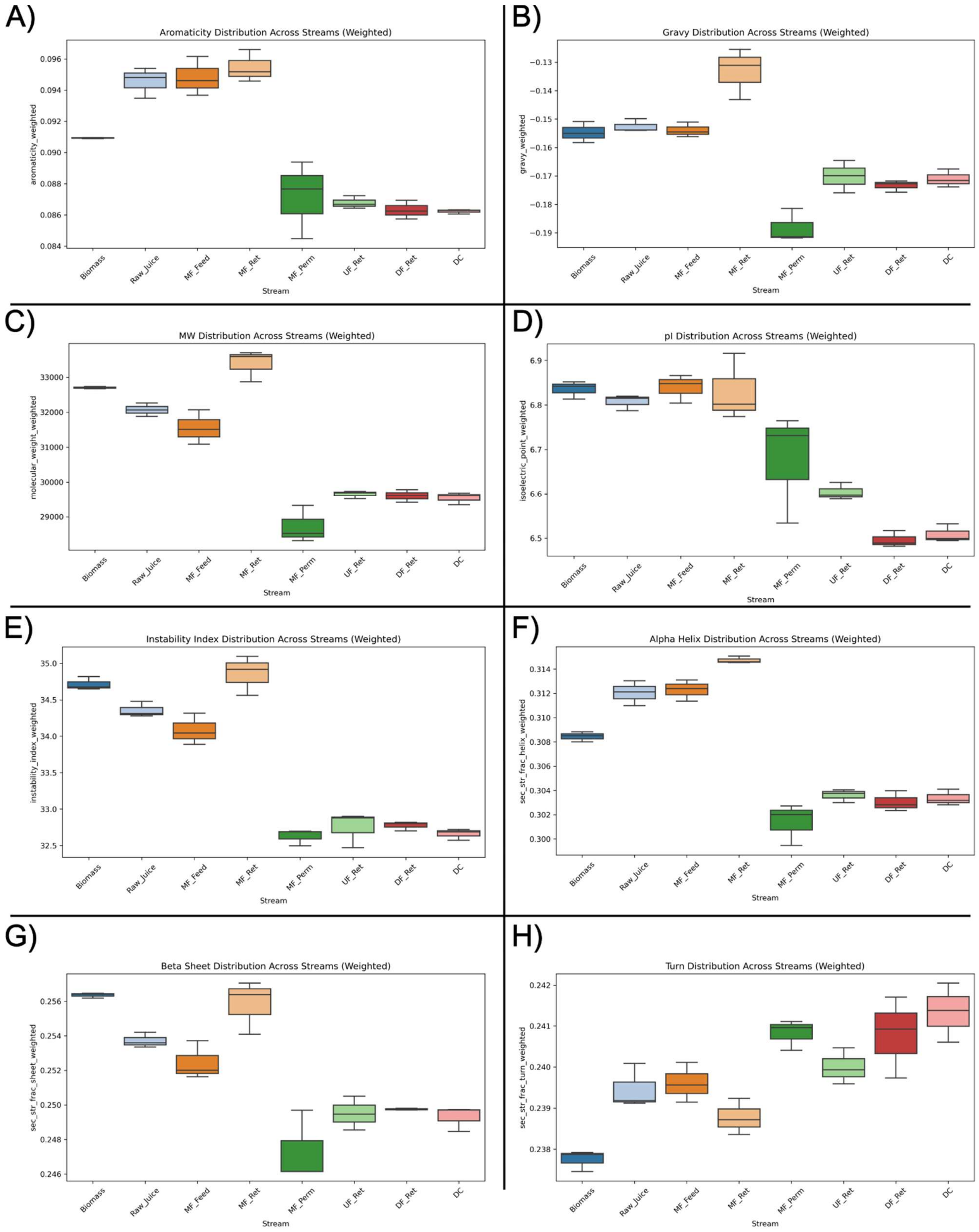


**Figure S9:** Boxplots showing the distribution of computed physicochemical properties by process stream (S2 Perm and DF Perm excluded) based on riBAQ-weighted analysis. Properties computed include: Aromaticity (A), GRAVY (B), molecular weight (C), isoelectric point (D), instability index (E), α-helical content (F), β-sheet content (G), and turn content (H).

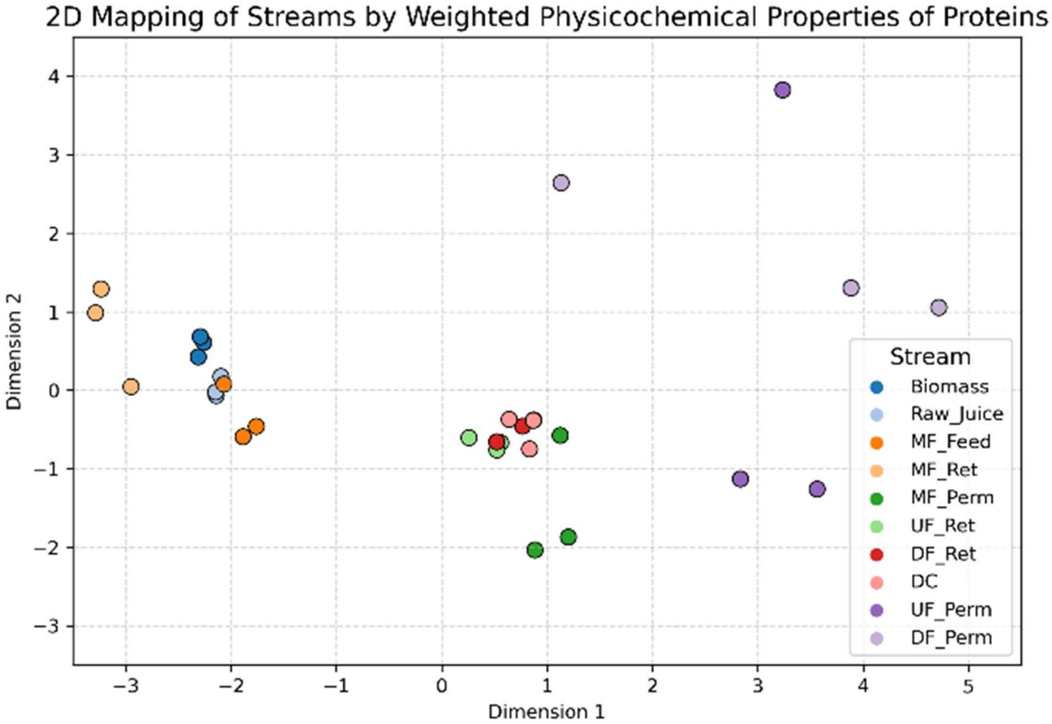


**Figure S10:** Two-dimensional representation of protein properties from each stream (replicate-level) obtained via TruncatedSVD on riBAQ-weighted computed physicochemical properties.
